# Evidence for nutrient-dependent regulation of the COPII coat by O-GlcNAcylation

**DOI:** 10.1101/2020.12.30.424839

**Authors:** Brittany J. Bisnett, Brett M. Condon, Noah A. Linhart, Caitlin H. Lamb, Duc T. Huynh, Jingyi Bai, Timothy J. Smith, Jimin Hu, George R. Georgiou, Michael Boyce

**Affiliations:** Department of Biochemistry, Duke University School of Medicine, Durham, NC 27710 USA

## Abstract

O-linked β-*N*-acetylglucosamine (O-GlcNAc) is a dynamic form of intracellular glycosylation common in animals, plants and other organisms. O-GlcNAcylation is essential in mammalian cells and is dysregulated in myriad human diseases, such as cancer, neurodegeneration and metabolic syndrome. Despite this pathophysiological significance, key aspects of O-GlcNAc signaling remain incompletely understood, including its impact on fundamental cell biological processes. Here, we investigate the role of O-GlcNAcylation in the coat protein II complex (COPII), a system universally conserved in eukaryotes that mediates anterograde vesicle trafficking from the endoplasmic reticulum. We identify new O-GlcNAcylation sites on Sec24C, Sec24D and Sec31A, core components of the COPII system, and provide evidence for potential nutrient-sensitive pathway regulation through site-specific glycosylation. Our work suggests a new connection between metabolism and trafficking through the conduit of COPII protein O-GlcNAcylation.

## Introduction

O-linked β-*N*-acetylglucosamine (O-GlcNAc) is a reversible monosaccharide modification of intracellular proteins, common in many clades of animals and plants (Bond and Hanover, 2015; Hanover et al., 2010; Hart, 2014; Hart et al., 2011). O-GlcNAc is added by O-GlcNAc transferase (OGT) and removed by O-GlcNAcase (OGA), often cycling on a timescale of minutes to effect cell signaling (Bond and Hanover, 2015; Hanover et al., 2010; Hart et al., 2011; King et al., 2019; Yang and Qian, 2017). O-GlcNAcylation is essential, as deletion of either OGT and OGA is lethal in mice (Keembiyehetty et al., 2015; Levine and Walker, 2016; Shafi et al., 2000; Yang et al., 2012), and O-GlcNAc signaling is dysregulated in a variety of diseases, including cancer, diabetes, cardiac dysfunction, neurodegeneration and X-linked intellectual disability (Akan et al., 2018; Bond and Hanover, 2013; Darley-Usmar et al., 2012; Dassanayaka and Jones, 2014; Erickson, 2014; Erickson et al., 2013; Hanover and Wang, 2013; Hardiville and Hart, 2014; Hart et al., 2011; Lazarus et al., 2009; Ma and Hart, 2013; Ma and Vosseller, 2013; Pravata et al., 2020a; Pravata et al., 2019; Pravata et al., 2020b; Selvan et al., 2018; Shin et al., 2011; Singh et al., 2015; Vaidyanathan et al., 2014; Vaidyanathan et al., 2017; Vaidyanathan and Wells, 2014; Wang and Hanover, 2013; Willems et al., 2017; Yi et al., 2012; Yuzwa et al., 2012; Yuzwa and Vocadlo, 2014; Zhu et al., 2014).

Despite this broad significance, important aspects of O-GlcNAcylation are incompletely understood. For example, for a handful of substrates, specific biochemical effects of O-GlcNAc have been documented, including conformational changes (Erickson et al., 2013; Tarrant et al., 2012), mediating or disrupting protein-protein interactions (Tarbet et al., 2018a; Tarbet et al., 2018b; Toleman et al., 2018) and altering protein stability (Han and Kudlow, 1997; Zhang et al., 2003) or subcellular location (Dentin et al., 2008). However, for the vast majority of substrates, the mechanistic effects of O-GlcNAc on its targets remain uncharacterized. Similarly, the cell-level signaling functions of O-GlcNAc are not fully elucidated. Of the thousands of O-GlcNAc substrates in mammals, many act in gene regulation or nutrient-sensing pathways, indicating that O-GlcNAcylation plays an essential role in those processes (Bond and Hanover, 2015; Hanover et al., 2010; Hart et al., 2011; King et al., 2019; Yang and Qian, 2017), but the key glycosylation events that govern these aspects of cell biology are not fully appreciated. New efforts are needed to understand the functions of O-GlcNAcylation in core cell biological processes.

We and others have previously reported extensive O-GlcNAcylation among the components of the coat protein II (COPII) complex (Boyce et al., 2011; Cho and Mook-Jung, 2018; Cox et al., 2018; Dudognon et al., 2004; Teo et al., 2010; Wells et al., 2002; Woo et al., 2015; Woo et al., 2018; Zachara et al., 2011; Zachara et al., 2004). COPII is the essential transport pathway that exports protein and lipid cargoes from the endoplasmic reticulum (ER) towards the Golgi in all eukaryotes (Aridor, 2018; Bethune and Wieland, 2018; Brandizzi, 2018; Gomez-Navarro and Miller, 2016; Hutchings and Zanetti, 2019; Peotter et al., 2019). Five proteins are necessary and sufficient for the formation of COPII carriers *in vitro:* The small GTPase Sar1, the inner coat proteins Sec23 and Sec24, and the outer coat proteins Sec13 and Sec31 (Aridor, 2018; Bethune and Wieland, 2018; Brandizzi, 2018; Gomez-Navarro and Miller, 2016; Hutchings and Zanetti, 2019; Matsuoka et al., 1998; Peotter et al., 2019). Through an ordered series of protein-protein interactions, COPII carriers assemble at ER exit sites, recruit protein cargoes into the vesicle, promote membrane curvature and scission, and mediate some aspects of target membrane tethering (Aridor, 2018; Bethune and Wieland, 2018; Brandizzi, 2018; Gomez-Navarro and Miller, 2016; Hutchings and Zanetti, 2019; Peotter et al., 2019). Although the minimal components of COPII vesicles are relatively well understood at the biochemical and structural levels, much less is known about how cells dynamically modulate COPII trafficking in response to stimuli or stress.

Post-translational modifications (PTMs), including O-GlcNAc, are emerging as one important mode of COPII regulation. Indirect evidence for a connection between O-GlcNAc and COPII has existed at least since 1987, when Lucocq and coworkers observed intracellular O-glycans of unknown origin marking ER exit sites in electron micrographs of porcine hepatocytes (Lucocq et al., 1987). More recently, we demonstrated that site-specific O-GlcNAcylation of Sec23A (one of two human Sec23 paralogs) is required for the trafficking of collagen, a large COPII client protein, in both human cells and developing zebrafish (Cox et al., 2018). Beyond Sec23, the cargo-binding protein Sec24 and the outer coat component Sec31 are also O-GlcNAc-modified (Boyce et al., 2011; Cho and Mook-Jung, 2018; Cox et al., 2018; Dudognon et al., 2004; Teo et al., 2010; Wells et al., 2002; Zachara et al., 2011; Zachara et al., 2004). However, the relevant upstream stimuli for, and downstream effects of, O-GlcNAcylation remain unknown.

Here, we use a combination of mass spectrometry (MS)-mediated O-GlcNAc site-mapping, CRISPR genome editing and biochemical and cell-based assays to investigate the functional impact of O-GlcNAcylation on Sec24 and Sec31 human proteins. These results provide evidence that COPII protein O-GlcNAcylation participates in both protein-protein interactions and nutrient-sensing, shedding new light on the regulation of trafficking in the early secretory pathway.

## Results

The outer COPII protein Sec31 is essential for carrier formation and is thought to be controlled by PTMs, including phosphorylation and ubiquitination (Hu et al., 2016; Jin et al., 2012; Koreishi et al., 2013; McGourty et al., 2016; Salama et al., 1997). We and others have also reported O-GlcNAcylation of Sec31A (Cho and Mook-Jung, 2018; Clark et al., 2008; Cox et al., 2018; Teo et al., 2010), one of two human paralogs, but the functional implications of this modification for the COPII system are still obscure. One possibility is that Sec31A is dynamically glycosylated as a regulatory function. To test this notion, we treated human cells with the OGT inhibitor Ac45SGlcNAc (5SGlcNAc) (Gloster et al., 2011) or the OGA inhibitor Thiamet-G (Yuzwa et al., 2008) and examined O-GlcNAcylation of endogenous Sec31A by immunoprecipitation and immunoblot (IP/IB) (Figure 1A). These results revealed that OGT inhibition decreased, and OGA inhibition increased, glycosylation of Sec31A, consistent with a regulatory, cycling (versus static) modification (Figure 1A).

**Figure 1.**
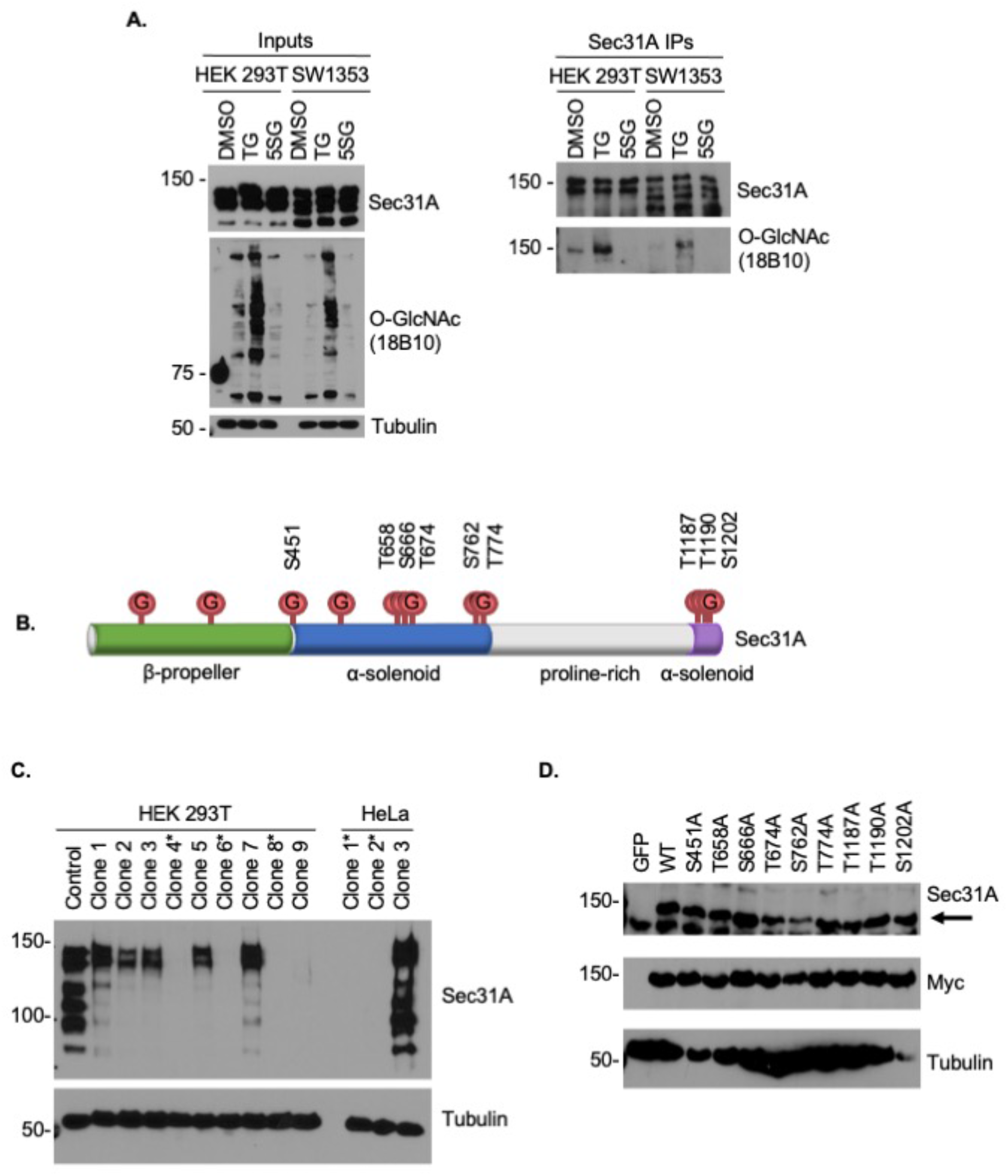
Sec31A is dynamically O-GlcNAcylated. **A.** Endogenous Sec31A was IP-ed from HEK 293T or SW1353 cells treated with DMSO, 50 μM Thiamet-G (TG), or 50 μM 5SGlcNAc (5SG) for 24 hours. Even Sec31A expression and IP efficiency were confirmed by Sec31A IB. Sec31A O-GlcNAc levels were detected using the anti-O-GlcNAc monoclonal antibody 18B10. **B.** ETD MS analysis indicates that human Sec31A contains at least 12 O-GlcNAcylated residues. Nine sites were previously detected, with resolution sufficient to localize O-GlcNAc on four residues: S451, T658, S666, and T674 (Cox et al., 2018). This study revealed two new sites (S762, T774) and allowed assignment of three previously ambiguous sites to T1187, T1190, and S1202. **C.** CRISPR-Cas9 methods were used to delete Sec31A from both HEK 293T and HeLa cells. Clonal cell lines created with a control or Sec31A-targeting sgRNAs were analyzed by IB. Asterisks indicate successful deletion of Sec31A. **D**. HEK 293T Sec31A^-/-^ clones were stably transduced with GFP, wild type (WT) Sec31A-myc6xHis, or Ser/Thr→Ala Sec31A-myc6xHis mutants corresponding to the nine O-GlcNAc sites identified in **B.** Expression of myc-6xHis-tagged Sec31A constructs in Sec31A^-/-^ cells was confirmed by Sec31A (arrow) and myc IB.

Like many O-GlcNAc substrates, COPII proteins are glycosylated on multiple residues (Cox et al., 2018). Our prior studies of Sec23A demonstrated that certain O-GlcNAc sites are essential for some of the trafficking functions tested thus far, whereas other sites are not (Cox et al., 2018). These results underline the importance of comprehensive site-mapping of O-GlcNAcylation in order to understand its impact on substrate proteins. We previously identified four O-GlcNAc sites on Sec31A (Cox et al., 2018), but these results were obtained with collision-induced dissociation mass spectrometry (CID MS), which is prone to false-negatives, due to the lability of the glycosidic bond in the CID fragmentation modality (Myers et al., 2013; Vercoutter-Edouart et al., 2015). Therefore, we hypothesized that additional human Sec31A glycosites may remain undiscovered. To address this knowledge gap, we performed O-GlcNAc site-mapping on Sec31A expressed in and purified from human cells using an MS method that includes electron transfer dissociation (ETD) MS (Figure 1B) (Vercoutter-Edouart et al., 2015). Indeed, ETD results pinpointed three O-GlcNAcylated residues, T1187, T1190, and S1202, that we had detected previously by CID but been unable to assign unambiguously to specific amino acids (Figure 1B) (Cox et al., 2018). In addition, ETD results revealed two new sites of glycosylation, S762 and T774, that have not been reported before (Figure 1B). Interestingly, all five of these newly assigned glycosites reside in the C-terminal half of Sec31A, in structural domains that participate in key protein-protein interactions within the COPII coat (Figure 1B) (Livak and Schmittgen, 2001; Ong et al., 2010; Stagg et al., 2008). These results suggest a potential influence of O-GlcNAc on the biochemical function of Sec31A and highlight the importance of thorough site-mapping in characterizing substrate O-GlcNAcylation.

Next, we sought to investigate the functional importance of individual Sec31A O-GlcNAc sites. Phenotypes from unglycosylatable point-mutants of O-GlcNAc substrates can be suppressed in the presence of the wild type protein, particularly for glycoproteins that function in multiprotein assemblies (Cox et al., 2018; Tarbet et al., 2018a). Therefore, null genetic backgrounds are critical systems for characterizing the effects of individual glycosites. To address this challenge, we used CRISPR methods to delete Sec31A from human cell lines and verified success by IB (Figure 1C). We then stably reexpressed epitope-tagged human wild type or individual selected unglycosylatable O-GlcNAc site mutants (S/T→A) in Sec31A^-/-^ cells, creating systems for further biochemical and functional characterization (Figure 1D). Some mutants exhibited lower expression, compared to wild type (Figure 1D). Because these mutants may express at lower levels due to technical variation in the protocol (e.g., position effects from gene insertion), we focused our subsequent experiments on the mutants expressed at wild type levels.

We used these reconstituted cells to identify the predominant glycosites on Sec31A in vehicle- and Thiamet-G-treated cells, to detect both constitutive and inducible sites (Figures 2A-B). We observed O-GlcNAcylation on all Sec31A variants, and Thiamet-G increased this glycosylation (Figures 2A-B), similar to our initial observations with endogenous protein (Figure 1A). However, specific point-mutants revealed differences. For example, compared to wild type, the S1202A mutant demonstrated consistently reduced O-GlcNAcylation in vehicle-treated cells (Figure 2B). This observation indicates that S1202 is a major Sec31A O-GlcNAc site under homeostatic conditions.

**Figure 2.**
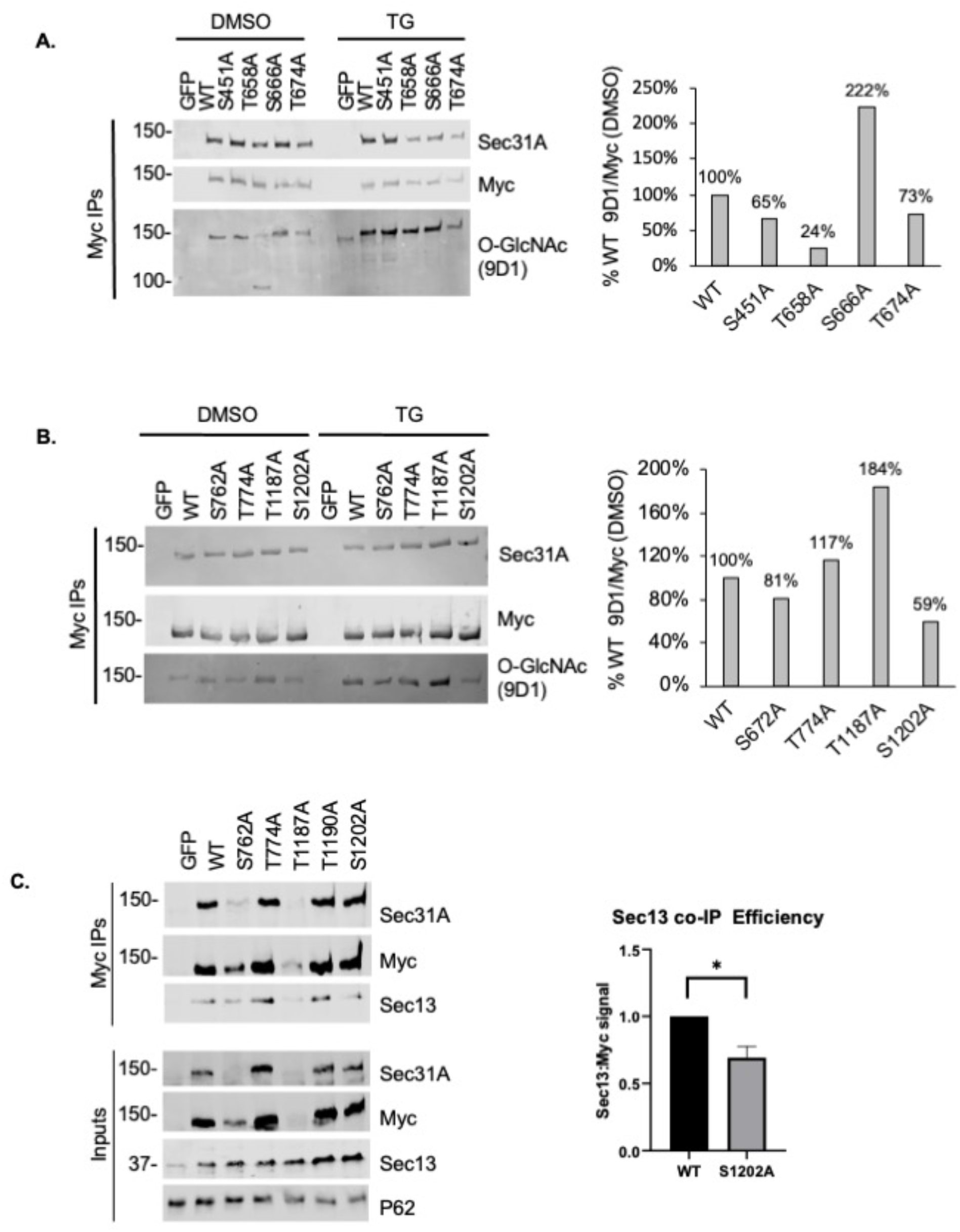
S1202 is a predominant O-GlcNAc site on Sec31A and is required for interaction with Sec13. **A.** and **B.** HEK 293T cells stably expressing GFP, myc-tagged WT or unglycosylatable Sec31A mutants were treated with DMSO or 50 μM TG for 24 hours. *Left:* Efficient IP of myc-6xHis-tagged WT Sec31A and Sec31A mutants was confirmed by quantitative IB for Sec31A and myc. Sec31A O-GlcNAcylation levels were measured using quantitative IB with anti-O-GlcNAc monoclonal antibody 9D1. *Right:* O-GlcNAc signal was normalized to myc signal for vehicle (DMSO)-treated samples and graphically displayed as a percentage of WT signal (representative blot). **C.** *Left:* HEK 293T cells stably expressing GFP, myc-6xHis-tagged WT or unglycosylatable Sec31A mutants were subjected to myc IP. Quantitative IBs were performed for Sec31A, myc, Sec13 and nucleoporin-62 (loading control). *Right:* Efficiency of Sec13 co-IP with WT or mutant Sec31A was assessed by dividing Sec13 signal by myc signal, normalized to WT value (n=4, * indicates *p* value < 0.05 by Student’s t-test).

O-GlcNAcylation can induce a range of biochemical effects on its substrates, including the modulation of protein-protein interactions (Tarbet et al., 2018b). Because the COPII coat functions through an intricate, highly orchestrated set of such interactions (Aridor, 2018; Bethune and Wieland, 2018; Brandizzi, 2018; Gomez-Navarro and Miller, 2016; Hutchings and Zanetti, 2019; Peotter et al., 2019), we tested whether the binding of Sec31A to known partners depends on individual O-GlcNAc sites. Notably, the interaction of Sec31A with its obligate outer coat binding partner, Sec13, was significantly reduced in the S1202A mutant, as compared to wild type (Figure 2C). These results suggest that site-specific O-GlcNAcylation influences specific protein-protein interactions in outer COPII coat assembly or disassembly.

Finally, as a first step towards dissecting the regulatory role of Sec31A O-GlcNAcylation, we investigated candidate upstream stimuli that might trigger glycosylation changes. We observed no consistent changes in Sec31A O-GlcNAcylation in response to calcium ionophore (Figure 3A), the ER stress inducer tunicamycin or rapamycin, an inhibitor of the nutrient-sensing kinases mTORC1/2 (Figure 3B), all stimuli that influence COPII trafficking in other contexts (Ge et al., 2017a; Helm et al., 2014; la Cour et al., 2013; Liu et al., 2019; McGourty et al., 2016). However, we observed a substantial increase in Sec31A O-GlcNAcylation in response to glucose reduction, as judged by multiple independent anti-O-GlcNAc monoclonal antibodies (Figure 3C). Taken together, these results indicate that nutrient-sensitive, site-specific O-GlcNAcylation of Sec31A may regulate its protein-protein interactions in the outer COPII coat and influence trafficking in the early secretory pathway. Efforts are underway to further test this hypothesis.

**Figure 3.**
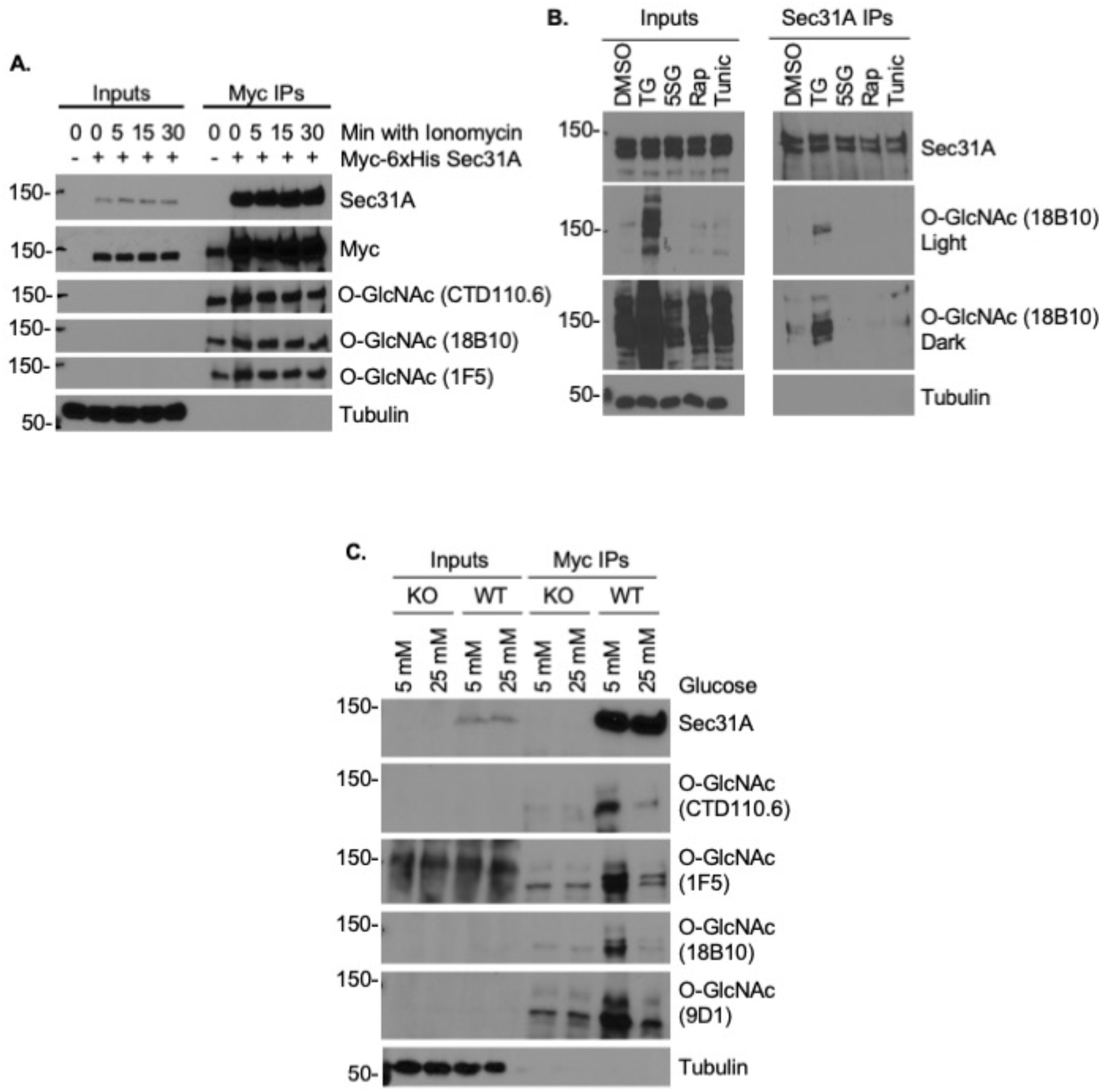
Lower glucose levels potentiate Sec31A O-GlcNAcylation. **A.** HEK 293T cells stably expressing GFP or WT Sec31A-myc-6xHis were treated with either DMSO or 3 μM ionomycin for 0, 5, 15, or 30 minutes. Sec31A was IP-ed (myc) and samples were analyzed by IB. CTD110.6, 18B10 and 9D1 are anti-O-GlcNAc monoclonal antibodies. **B.** HEK 293T cells were treated with DMSO, 50 μM TG or 50 μM 5SG for 24 hours, 200 nM rapamycin (Rap) for 16 hours, or 2.5 μg/mL tunicamycin (Tunic) for 5 hours. Endogenous Sec31A was IP-ed and samples were analyzed by IB. **C**. Sec31A^-/-^ HEK 293T cells stably expressing GFP (KO) or Sec31A-myc6xHis (WT) were grown in 5 mM or 25 mM glucose-containing medium for 24 hours. Samples were analyzed by IP/IB.

Similar to Sec31A, the inner coat proteins Sec23 and Sec24 are extensively O-GlcNAcylated in mammalian cells (Boyce et al., 2011; Cox et al., 2018; Dudognon et al., 2004; Lee et al., 2016; Teo et al., 2010; Woo et al., 2015; Woo et al., 2018; Zachara et al., 2011). We have previously shown that specific O-GlcNAcylation sites on Sec23A are required for endogenous collagen trafficking in cultured human cells and live zebrafish (Cox et al., 2018). However, the functional impact of O-GlcNAcylation on Sec24 remains unclear. Of the four human Sec24 paralogs, we first focused on Sec24C because we and other groups have confirmed it as a robust OGT substrate, and a prior report suggested a regulatory function for Sec24C O-GlcNAcylation (Boyce et al., 2011; Cox et al., 2018; Dudognon et al., 2004). To test the hypothesis that dynamic glycosylation governs Sec24C function, we used an IP/IB assay to confirm that endogenous human Sec24C is O-GlcNAcylated, with 5SGlcNAc treatment decreasing glycosylation, and Thiamet-G increasing it (Figures 4A-B). We CRISPR-deleted Sec24C in human cells (Figure S1) and repeated this experiment, confirming the specificity of the Sec24C and O-GlcNAc antibodies for Sec24C glycosylation (Figure 4A-B). Sec24C glycosylation is remarkably dynamic, as O-GlcNAcylation increased in as little as five minutes upon treatment with Thiamet-G and peaked at 8 hours, as judged by multiple anti-O-GlcNAc antibodies (Figure 4C-E). This result is consistent with a regulatory, rather than structural or static, function of Sec24C glycosylation.

**Figure 4.**
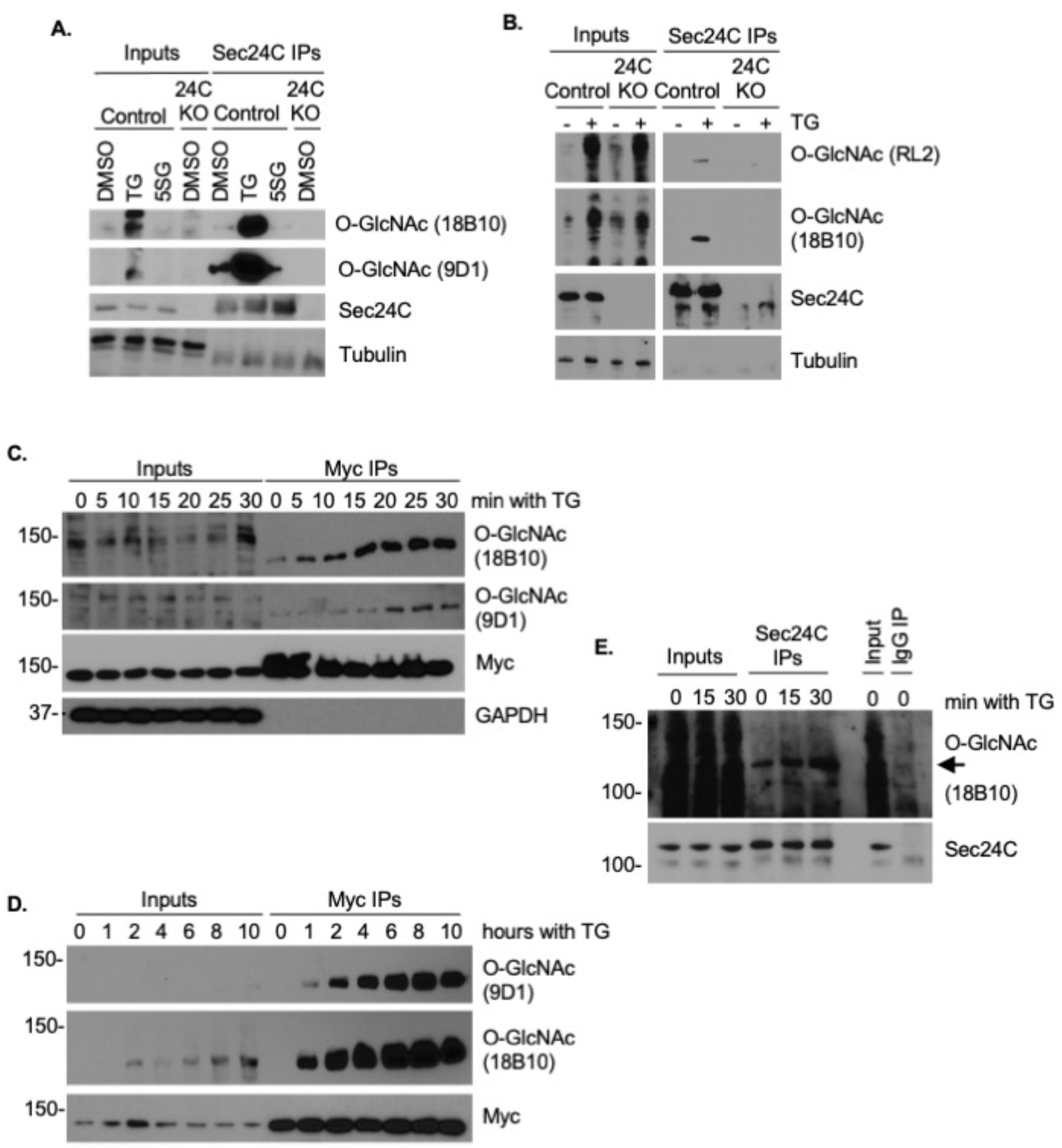
Sec24C is dynamically O-GlcNAcylated. **A.** Endogenous Sec24C was IP-ed from HEK 293T cells or Sec24C^-/-^ (KO) HEK 293T cells with treated with 50 μM TG, 5SG or vehicle control (DMSO) for 6 hours. An increase in Sec24C O-GlcNAcylation was detected with two O-GlcNAc monoclonal antibodies by IB. **B.** Endogenous Sec24C was IP-ed from control or Sec24C^-/-^ HeLa cells with treated with 50 μM TG or vehicle control for 6 hours. An increase in Sec24C O-GlcNAcylation with TG was detected with two O-GlcNAc monoclonal antibodies via IB. **C.** HEK 293T cells stably expressing myc-6xHis-tagged Sec24C were treated with 25 μM TG for 0-30 min. Cells were collected at 5-minute intervals and Sec24C O-GlcNAcylation was observed by myc IP and IB. **D.** HEK 293T cells stably expressing myc-6xHis-tagged Sec24C were treated with 25 μM TG for 0-10 hours. Cells were collected at 1, 2, 4, 8 and 10 hours post-treatment and Sec24C O-GlcNAcylation was analyzed by myc IP and IB. **E.** Endogenous Sec24C O-GlcNAcylation was analyzed by IP/IB from HEK 293T cells treated with 50 μM TG for 0, 15 or 30 min.

To further evaluate the significance of Sec24C O-GlcNAcylation, we performed ETD MS site-mapping, as above (Figure 5A-B). These results identified four previously unreported O-GlcNAcylation sites in the N-terminal low-complexity domain of Sec24C, more than doubling the number of glycosites that we identified previously with CID MS (Figure 5A-B) (Cox et al., 2018). To investigate the role of individual glycosylation sites, we re-expressed wild type or unglycosylatable mutant (S/T→A) cells in Sec24C^-/-^ cells (Figure 5C). Next, we assessed the potential importance of particular Sec24C glycosites. Using an IP/IB assay in both control and Thiamet-G-treated cells, we found that several mutants displayed significantly decreased O-GlcNAcylation, including S66A, S72A, S191A, T775A and T776A, and particularly in Thiamet-G-treated samples (Figure 6A-B). Interestingly, the S72A mutant also exhibited a higher degree of Sec23A binding than did wild type Sec24C under vehicle-treated conditions, perhaps indicating that constitutive O-GlcNAcylation at this site reduces this interaction (Figure 6A). These observations suggest that these five residues may be predominant sites of inducible O-GlcNAcylation and sites of Sec24C regulation.

**Figure 5.**
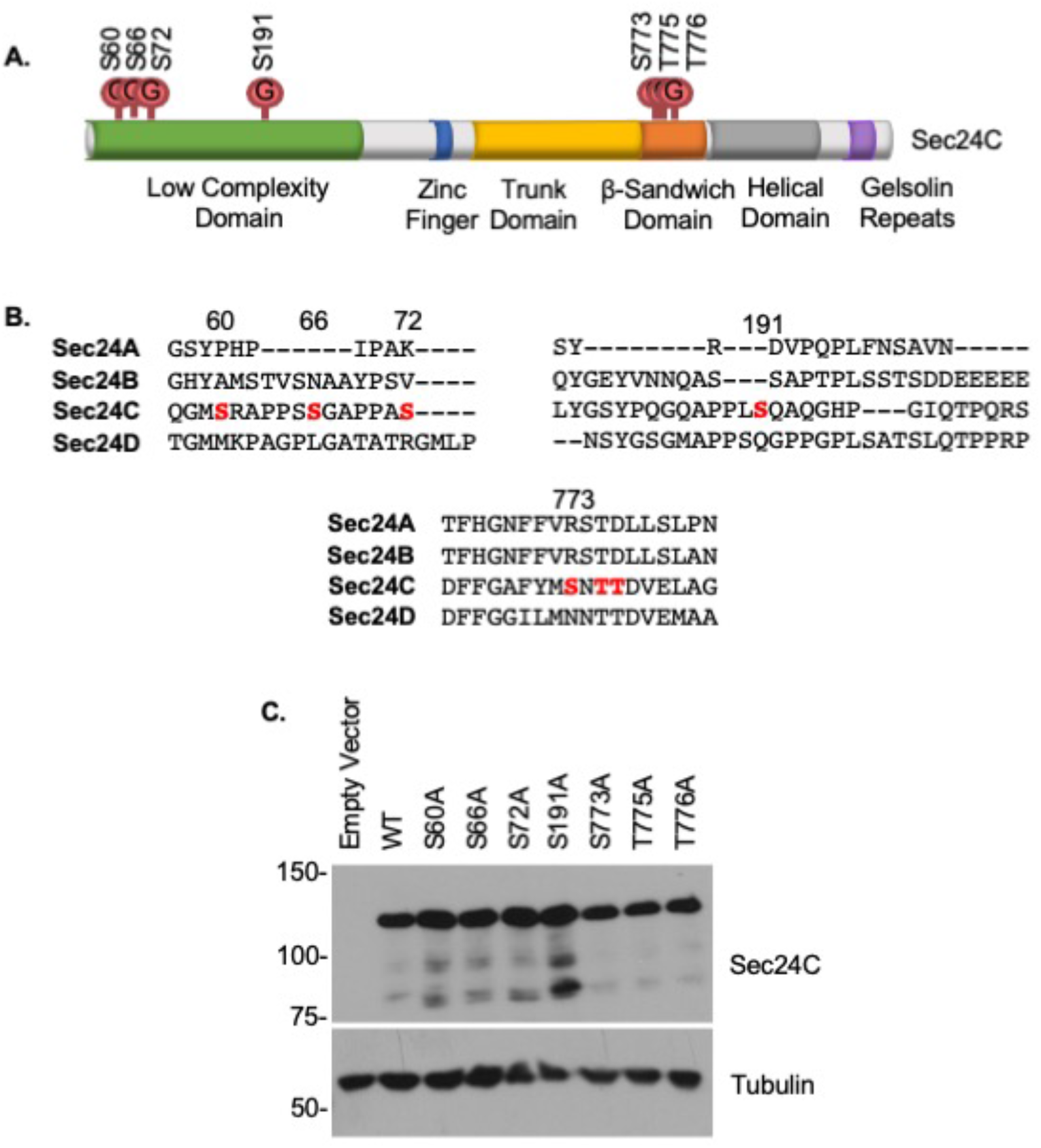
ETD MS reveals four new sites of Sec24C O-GlcNAcylation. **A.** Human Sec24C contains at least seven O-GlcNAc sites. S773, T775 and T776 were previously reported (Cox et al., 2018). Four additional sites were identified in this work via ETD MS in the N-terminal low complexity domain: S60, S66, S72 and S191. **B.** Conservation of Sec24C O-GlcNAc sites (red) among human Sec24 paralogs. **C.** IB verification of unglycosylatable Sec24C mutant expression in Sec24C^-/-^ HEK 293T cells.

**Figure 6.**
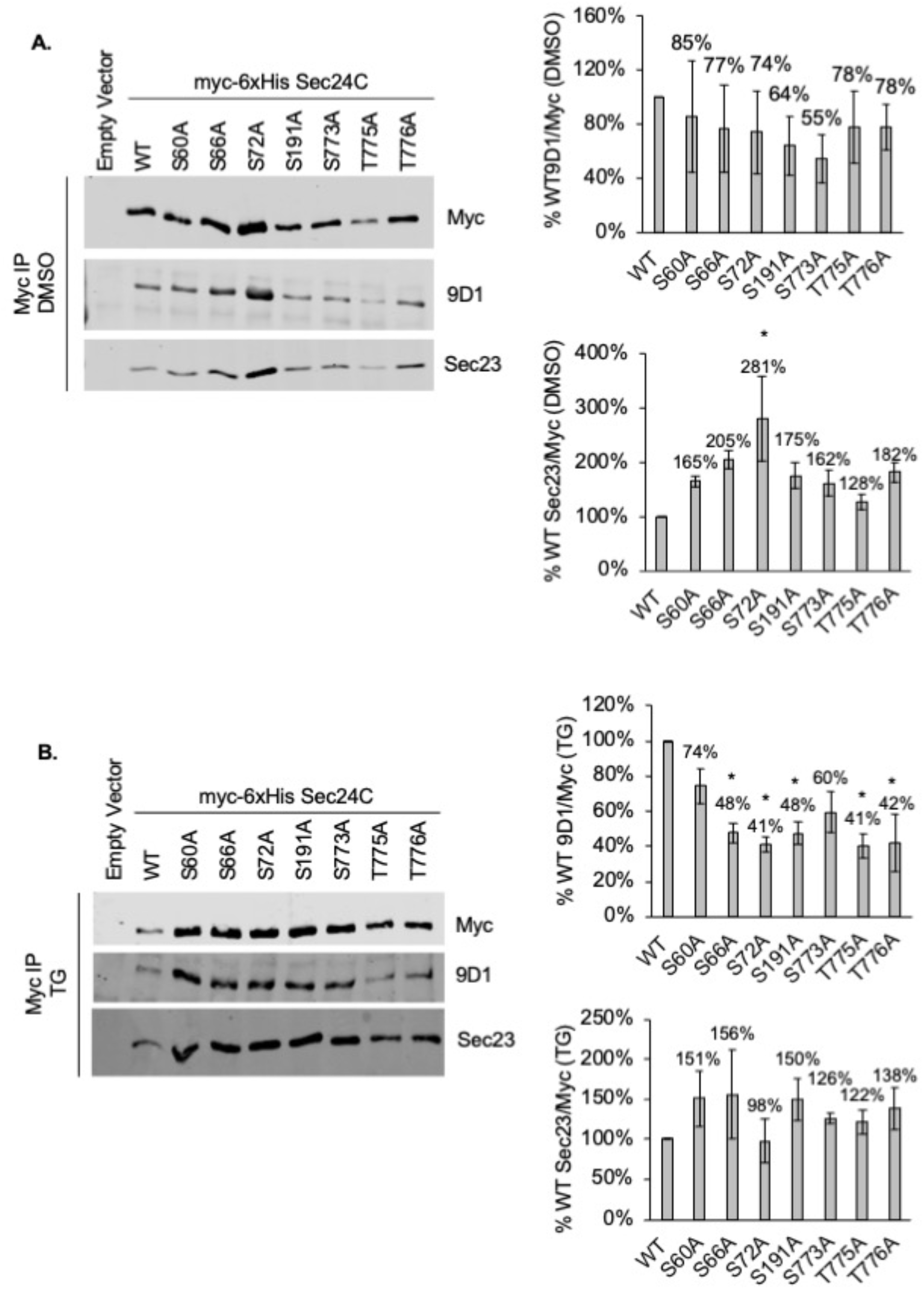
Identification of constitutive and inducible Sec24C O-GlcNAc sites. HEK 293T Sec24C^-/-^ cells expressing unglycosylatable mutants were treated with DMSO vehicle (**A**.) or 50 μM Thiamet-G (**B.**) for 6 hours and samples were analyzed by myc IP and quantitative IBs. Quantification and statistics were performed on three biological replicates. Error bars represent standard error of the mean (SEM). Asterisk indicates *p* < 0.05, compared to wild type, as determined by ANOVA and Tukey’s-HSD.

We also examined potential upstream stimuli that might influence Sec24C glycosylation. As with Sec31A, calcium ionophore produced no detectable changes on Sec24C O-GlcNAcylation (Figure S2). By contrast, we observed a dose-dependent increase in Sec24C O-GlcNAcylation in response to rapamycin (Figure 7A). This result suggests that Sec24C glycosylation may also be nutrient-responsive, and could be reciprocally regulated either by mTORC1 itself or a downstream kinase, such as Akt, which is a direct mTORC1 target and has been reported to phosphorylate Sec24C (Sharpe et al., 2011). mTORC1 is well known to regulate autophagy, and previous reports have indicated a functional role for Sec24C and other COPII components in contributing membrane to autophagosome biogenesis (Carlos Martín Zoppino et al., 2010; Ge et al., 2017b; Ge et al., 2014; Ishihara et al., 2001; Karanasios et al., 2016; Kim and Guan, 2019). Indeed, we observed an increase in the cleavage of the autophagosome marker LC3 in Sec24C^-/-^ cells (Figure 7B), implying an alteration in autophagy under homeostatic conditions. Together, these results suggest that Sec24C O-GlcNAcylation may serve a signaling function in autophagic flux, an important hypothesis to test in future work.

**Figure 7.**
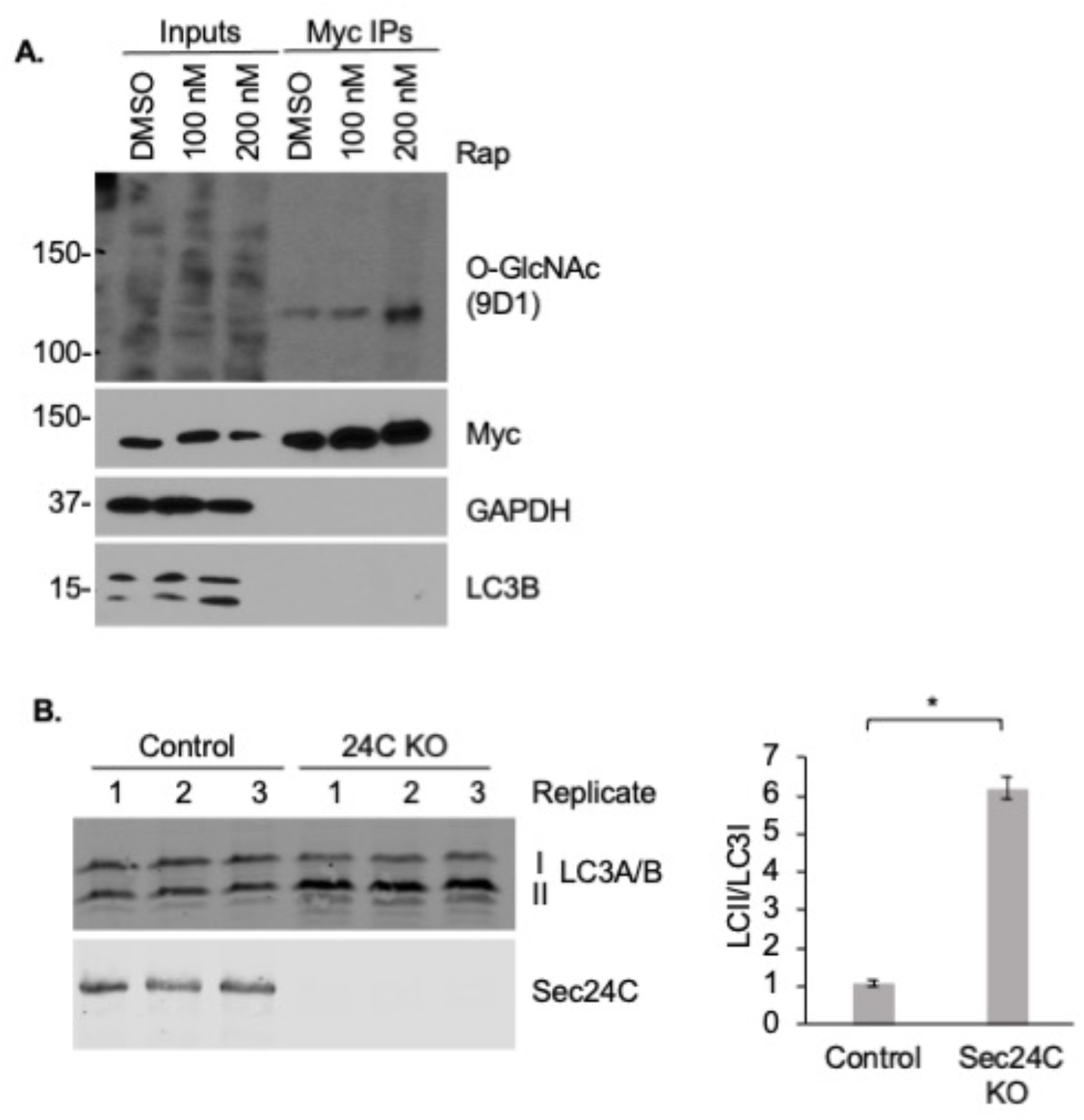
Rapamycin treatment increases Sec24C O-GlcNAcylation. **A.** HEK 293T cells stably expressing myc-6xHis Sec24C were treated with 100 nM or 200 nM rapamycin (Rap) or vehicle control for 16 hours. Sec24C was analyzed by myc IP and anti-O-GlcNAc IB (n=3). **B.** IB revealed that Sec24C^-/-^ (KO) HeLa cells have a higher ratio of LC3II/LC3I than do control cells (n=3). Error bars represent SEM. Asterisk indicates *p* < 0.05 as determined by Student’s t-test.

Finally, we investigated whether O-GlcNAcylation might be a broader mode of Sec24 regulation. Humans, mice and other mammals express four Sec24 paralogs, frequently in the same cell or tissue type, but the utility of producing multiple isoforms is not fully understood. One possibility is that PTMs may affect each paralog differently, affording finer combinatorial control of the COPII pathway. To test whether O-GlcNAcylation could, in principle, participate in such signaling, we examined Sec24D as another model paralog. Indeed, IP/IB experiments confirmed that 5SGlcNAc reduced, and Thiamet-G potentiated, the O-GlcNAcylation of epitope-tagged human Sec24D (Figure 8A). Furthermore, IP/IB revealed that endogenous Sec24D is also glycosylated, as confirmed using CRISPR-generated Sec24D^-/-^ cells as a specificity control (Figures 8B and S3). We used ETD MS to site-map O-GlcNAc on Sec24D and discovered six glycosites clustered in two discrete regions of the protein (Figures 8C). Notably, three of these reside in the N-terminal low-complexity domain (Figures 8C), reminiscent of our observations for Sec24C (Figure 5A). Together, these results indicate that O-GlcNAcylation may be a widely used mode of Sec24 regulation in mammalian cells.

**Figure 8.**
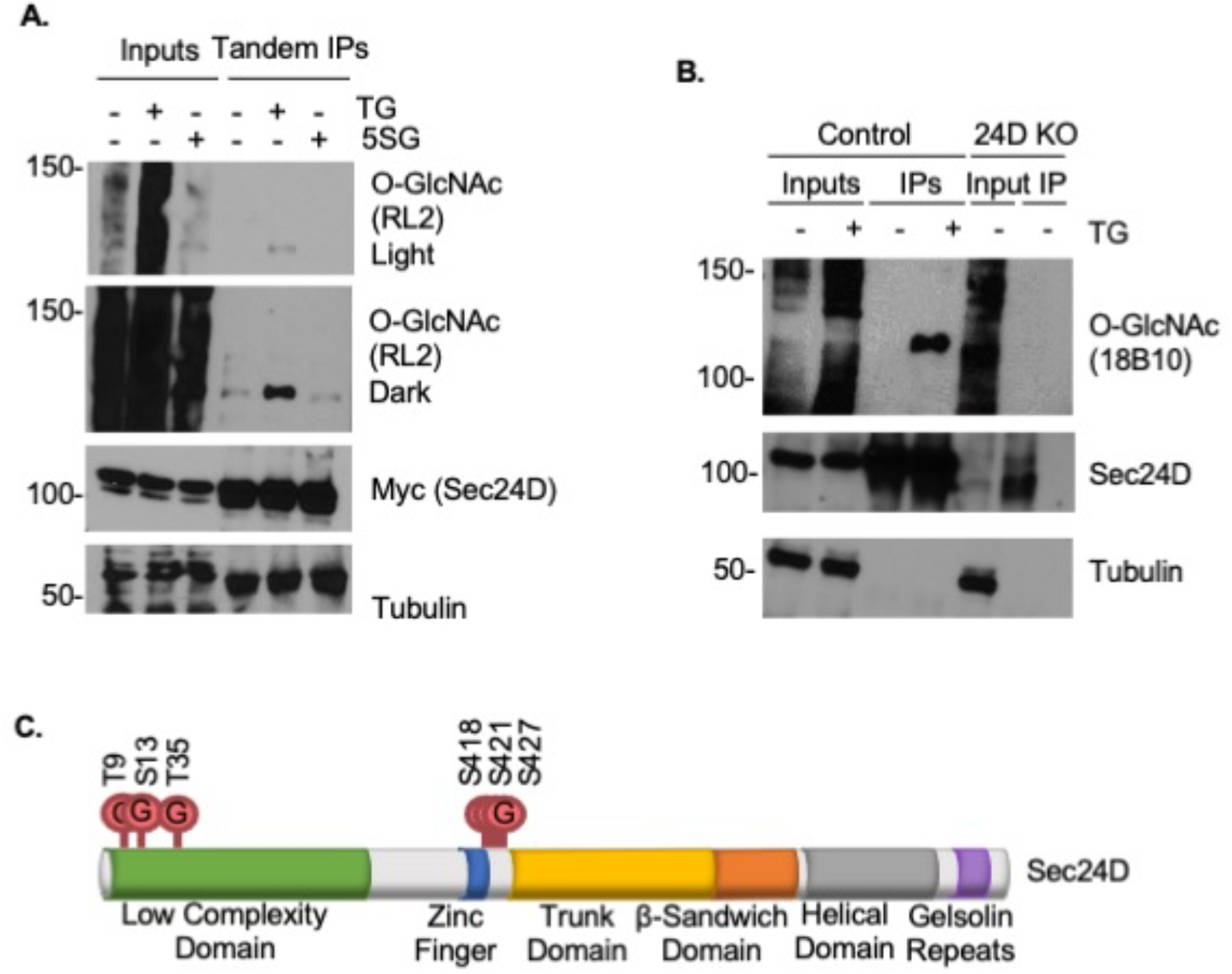
Sec24D is dynamically O-GlcNAcylated. **A.** Myc-6xHis Sec24D was transiently expressed in HEK 293T cells treated with 50 μM TG, 5SG, or DMSO vehicle control for 6 hours. Sec24D was analyzed by tandem myc IP/Ni-NTA affinity purification and O-GlcNAc IB. **B.** Endogenous Sec24D was IP-ed from control Sec24D^-/-^ (KO) HEK 293T cells treated with 50 μM TG or DMSO vehicle control for 6 hours and analyzed by O-GlcNAc IB. **C.** Six O-GlcNAc sites were identified on Sec24D by ETD MS: T9, S13, T35, S418, S421, T427.

## Discussion

O-GlcNAc is a ubiquitous and essential modification in nearly all mammalian cells examined. However, the biochemical effects that O-GlcNAc has on its substrates and its impact on downstream processes in mammalian cells remain incompletely understood. Here, we have addressed these questions in the context of the COPII trafficking pathway. Decades of elegant biochemical, genetic and structural studies have illuminated the mechanisms and functions of the core COPII proteins (Aridor, 2018; Barlowe, 2020; Bethune and Wieland, 2018; Brandizzi, 2018; Gomez-Navarro and Miller, 2016; Hutchings and Zanetti, 2019; Peotter et al., 2019). By contrast, much less is known about how mammalian cells adjust COPII activity in response to fluctuating signals or stresses. Taken together, our results provide evidence for the overarching hypothesis that nutrient-sensitive O-GlcNAcylation of COPII proteins regulates vesicle trafficking.

Our studies on the outer coat protein Sec31A reveal extensive O-GlcNAcylation, including in the C-terminal domain (Figure 1B). This region of the protein is thought to be largely disordered, and yet specific motifs have been identified within it that mediate crucial functional interactions with other COPII proteins, including Sec23 and Sar1 (Bi et al., 2007; Fath et al., 2007; Stancheva et al., 2020). Recent reports suggest that the unstructured domain of Sec31 serves as a multivalent interface, engaging in numerous weak interactions with other COPII components to confer sufficient rigidity on the outer coat cage, while also allowing for ready reversibility (e.g., through competing interactions with proteins on the target membrane) (Hutchings et al., 2018; Hutchings and Zanetti, 2019; Markova and Zanetti, 2019). It is interesting to speculate that PTMs occurring in this C-terminal domain, such as O-GlcNAc (Figure 1B), may serve to inhibit or potentiate some of these multivalent interactions, providing finer control over COPII carrier geometry, uncoating kinetics or protein-protein interactions.

In particular, we demonstrate that S1202, a predominant O-GlcNAc site (Figure 2A-B), is required for wild type levels of Sec13 binding (Figure 2C), suggesting that O-GlcNAcylation might modulate vesicle trafficking through the Sec31A/Sec13 protein-protein interaction. This hypothesis is consistent with the general model of COPII vesicle formation as a precisely choreographed sequence of subunit binding events (Aridor, 2018; Bethune and Wieland, 2018; Brandizzi, 2018; Gomez-Navarro and Miller, 2016; Hutchings and Zanetti, 2019; Peotter et al., 2019), and with a substantial body of literature showing that O-GlcNAc often acts by altering protein-protein interactions (Tarbet et al., 2018b). However, in other ways, these results are surprising. For example, the known Sec13 binding site resides at the Sec31 N-terminus, distant in primary sequence space from S1202 (Fath et al., 2007; Yamasaki et al., 2006). The disordered C-terminal region of Sec31 may exhibit considerable flexibility *in vivo*, influencing Sec13 binding in unexpected ways. Indeed, the best available structural information on Sec31 has been obtained with the *S. cerevisiae* ortholog (Fath et al., 2007; Lederkremer et al., 2001; Matsuoka et al., 2001; Stagg et al., 2006; Stancheva et al., 2020), which has relatively modest homology (27% amino acid identity, 47% similarity) to the human protein, leaving open the possibility that the human Sec31A structure may be considerably different. Alternatively, it may be that O-GlcNAcylation at S1202 induces a conformational or other allosteric change that affects Sec13 binding at another site. Finally, we note that a prior study also reported several O-GlcNAc sites in the C-terminal domain of Sec31A, distinct from the ones we identified here, and suggested a role for these putative modifications in mediating proteinprotein interactions (Cho and Mook-Jung, 2018). However, this report relied almost exclusively on indirect affinity purifications with wheat germ agglutinin, a promiscuous lectin, and did not provide any direct evidence of Sec31A O-GlcNAcylation (e.g., MS site-mapping or quantitative IB of purified wild type and mutant protein) (Cho and Mook-Jung, 2018). Therefore, O-GlcNAcylation at these reported sites remains unproven.

We also investigated the potential signaling relevance of Sec31A O-GlcNAcylation. It has long been known that COPII vesicle formation can be reconstituted *in vitro* without native PTMs (Matsuoka et al., 1998). This result indicates that modifications of Sec31A and other coat components are likely not required for carrier formation *in vivo*, but instead may serve a regulatory function, such as tuning the pathway in response to stimuli or stress. Consistent with this idea, we found that glucose reduction increased Sec31A O-GlcNAcylation (Figure 3C). Uridine diphosphate-GlcNAc (UDP-GlcNAc), the nucleotide-sugar cofactor for OGT, is the end-product of the hexosamine biosynthetic pathway (HBP), which consumes approximately 2-5% of glucose in mammalian cells (Hanover et al., 2010). Because a range of metabolites feeds into the HBP, UDP-GlcNAc levels and O-GlcNAcylation are often considered sentinels of nutrient availability, fluctuating as metabolite levels rise and fall (Akan et al., 2018; Bond and Hanover, 2013; Hanover et al., 2010). In the case of Sec31A, it may initially appear counter-intuitive that a decrease in glucose results in an increase in O-GlcNAcylation, but analogous observations have been reported previously for OGT substrates in other contexts (Ruan et al., 2017; Taylor et al., 2009; Zou et al., 2012). Prior research has also demonstrated changes in COPII trafficking in response to the availability of glucose and other nutrients, but the underlying molecular mechanisms are not fully understood (Amodio et al., 2009; Jeong et al., 2018; Liu et al., 2019; Zacharogianni et al., 2014). We speculate that glucose-responsive Sec31A O-GlcNAcylation may serve as one link between nutrient-sensing and COPII function, perhaps adjusting the trafficking of cargoes such as growth factor receptors or metabolite transporters in response to glucose availability. Future work will test these hypotheses directly.

Our results indicate a potential regulatory role for O-GlcNAcylation on the Sec24 inner coat proteins as well. Consistent with earlier reports (Boyce et al., 2011; Cox et al., 2018; Dudognon et al., 2004; Teo et al., 2010; Woo et al., 2015; Woo et al., 2018; Zachara et al., 2011), our MS site-mapping results revealed extensive glycosylation of Sec24C, including previously undiscovered sites (Figures 4–5). Remarkably, O-GlcNAc cycles very rapidly on Sec24C, as revealed by the clear increase in glycosylation within five minutes of applying an OGA inhibitor (Figure 4C,E). These observations suggest the intriguing hypothesis that O-GlcNAc may be added to and removed from Sec24C (perhaps more than once) within a single cycle of COPII vesicle forming, trafficking, uncoating and fusing with its target membrane. Previous reports have suggested a role for temporally or spatially restricted modification of COPII components, such as phosphorylation of Sec23 by the Golgi-tethered kinase Hrr25p/CKIδ, which is proposed to control vesicle uncoating and enforce unidirectional trafficking (Bhandari et al., 2013; Lord et al., 2011). Future experiments, such as proximity ligation strategies, will use imaging methods to visualize the glycosylated and total pools of Sec24C in fixed cells and determine whether spatially restricted O-GlcNAcylation impacts on Sec24 and COPII function.

Our data also lend new support to prior proposals that Sec24C is subject to reciprocal regulation by O-GlcNAc and phosphorylation (Dudognon et al., 2004). The Paccaud group reported that Sec24C phosphorylation and O-GlcNAcylation appear to be mutually exclusive, with the former occurring in mitosis and the latter in interphase, although the relevant O-GlcNAc sites and kinase(s) were not determined (Dudognon et al., 2004). More recently, Akt was reported to phosphorylate Sec24C (Sharpe et al., 2011), and our results demonstrate that rapamycin, which inhibits mTORC complexes, induces Sec24C O-GlcNAcylation (Figure 7A). mTORC complexes directly phosphorylate Akt, potentiating its activity (Bhaskar and Hay, 2007). Together, these results suggest possible reciprocal modification by OGT and Akt, mTORC or other kinases. In the case of Akt, O-phosphate and O-GlcNAc may occur on distinct sites, as Akt reportedly phosphorylates Sec24C on S888 (Sharpe et al., 2011), which was not among the glycosites we detected (Figure 5A). However, reciprocal modification by O-phosphate and O-GlcNAc at distinct sites has been observed previously, including on Akt itself (Butkinaree et al., 2010; Leney et al., 2017; Wang et al., 2012; Zhong et al., 2015). Conversely, among the Sec24C glycosites we identified, T776 was reported to be phosphorylated in a study of mitotic substrates of Aurora and Pololike kinases (Kettenbach et al., 2011), potentially explaining the original Paccaud observation. Further experiments will be required to dissect the Sec24C sites, the upstream kinases and the downstream effects of this putative reciprocal regulation.

Interestingly, our results may indicate a role for Sec24C O-GlcNAcylation in nutrient-sensing as well. mTORC complexes are well-known regulators of autophagy, and rapamycin inhibition mimics a starvation signal, inducing autophagy (Al-Bari, 2020; Condon and Sabatini, 2019; Saxton and Sabatini, 2017). O-GlcNAcylation in general has been implicated in autophagy in a number of ways (Akan et al., 2018; Fahie and Zachara, 2016), and Sec24 in particular is reported to participate in contributing membrane to the nascent autophagosome in both macroautophagy and ER-phagy (Cui et al., 2019; Davis et al., 2016; Graef et al., 2013). Consistent with these studies, we showed that cleavage of the autophagy marker LC3 is increased at baseline in Sec24C^-/-^ cells (Figure 7B), indicating a dysregulation of autophagic flux. In the future, it will be important to determine whether site-specific Sec24C glycosylation influences autophagosome formation, and whether mTORC signaling impacts on this regulation.

Lastly, we provide evidence that O-GlcNAcylation is likely a conserved mode of regulation among Sec24 paralogs. Mammals express four Sec24 paralogs, A through D, and the phenotypes of the individual Sec24 mouse knockouts are strikingly divergent (Adams et al., 2014; Baines et al., 2013; Chen et al., 2013; Merte et al., 2010; Wang et al., 2018; Wansleeben et al., 2010). However, all paralogs are thought to play similar or identical biochemical roles in the COPII inner coat, and recent evidence demonstrates that ectopic Sec24D expression can rescue the developmental defects in the nervous system of Sec24C^-/-^ mice (Wang et al., 2018). These results argue that Sec24 paralogs serve at least partially overlapping biochemical functions, and that tissue-specific expression patterns may largely explain the discrepancies in knockout phenotypes. If this model is correct, however, the advantage of expressing four paralogous Sec24 proteins is unclear. One possibility is that Sec24 isoforms may be differentially subjected to regulation by PTMs, providing another layer of combinatorial tuning of COPII activity. We identified six new sites on Sec24D, supporting the notion that O-GlcNAc is a common feature among Sec24 paralogs (Figure 8C). Three of these lie in the low-complexity domain, reminiscent of our observations with Sec24C, but at sites distinct from where Sec24C is modified, potentially implying differential regulation (Figure 5A). The functional significance of the low-complexity domains and their PTMs remain unclear, but O-GlcNAc moieties in this region may influence protein-protein interactions and COPII coat assembly or disassembly. Experiments are underway to test the potential requirement for specific Sec24C and Sec24D O-GlcNAc sites in vertebrate development, using rescue experiments in loss-of-function zebrafish models (Sarmah et al., 2010).

In sum, we have used the model proteins Sec24C and Sec31A to provide evidence that sitespecific O-GlcNAcylation may form a link between cellular metabolism and COPII trafficking. It has long been known that O-GlcNAc serves a nutrient-sensing function in other contexts (Bond and Hanover, 2015; Hanover et al., 2010; Hart et al., 2011; King et al., 2019; Yang and Qian, 2017). Conversely, COPII trafficking is appreciated to be dynamically regulated by both physiological stimuli and stresses, but the underlying molecular mechanisms are not fully understood. Our work suggests that the well-described nutrient-sensing function of O-GlcNAc may be brought to bear on COPII trafficking, with downstream consequences for protein secretion and function. Future work will determine the specific roles of sitespecific Sec24 and Sec31 glycosylation in cell and tissue physiology.

## Materials and methods

### Chemical synthesis

Ac_4_5SGlcNAc (abbreviated 5SG in figures) was synthesized as described (Gloster et al., 2011) and was a gift of Dr. Benjamin M. Swarts (Central Michigan University). Thiamet-G (abbreviated TG in figures) was synthesized as described (Yuzwa et al., 2008) by the Duke Small Molecule Synthesis Facility. All other chemicals were purchased from Sigma-Aldrich unless otherwise indicated.

### Cell culture

SW1353, HEK 293T and HeLa cell lines (as well as their engineered derivatives) were grown at 37 °C and 5% CO_2_ in Dulbecco’s Modified Eagle’s medium containing 10% fetal bovine serum, 100 units/mL penicillin, and 100 μg/mL streptomycin. Sec31A^-/-^ HEK 293T cell lines stably expressing eGFP, wild type Sec31A or unglycosylatable Sec31A mutants were supplemented with 200 μg/mL Zeocin.

### Mammalian expression vectors

pSpCas9(BB)-2A-GFP (PX458) was a gift from Dr. Feng Zhang (Addgene plasmid # 48138; http://n2t.net/addgene:48138; RRID:Addgene_4 8138) (Ran et al., 2013). The following single guide RNAs (sgRNAs) targeting *sec24C, sec24D*, and *sec31A* were designed and validated via the Surveyor assay (Ran et al., 2013) by the Duke Functional Genomics Facility:

sgRNA 24C-1: 5’ – TGCCCAAATGGTGGCACAGG – 3’
sgRNA 24C-2: 5’ – TATCATCAGTCCAGCTATGG– 3’
sgRNA 24D-1: 5’ – CCATAGTGCCCATAATGAGG – 3’
sgRNA 24D-2: 5’ – GGCTGAGGCTGAGAATACGG – 3’
sgRNA 24D-3: 5’ – ATTTTCATAATGAGTCAACA – 3’
sgRNA 31A-1: 5’ – CTTTAACTTCATCCTGCTAA – 3’
sgRNA 31A-2: 5’ – CTTTAGCAGGATGAAGTTAA – 3’
sgRNA 31A-3: 5’ – AACCTCATACCTGTGAGAAG – 3’

An sgRNA targeting the *AAVS1* “safe harbor” locus was used as a control (Sadelain et al., 2011). pLenti CMV/TO Puro DEST (670-1) was a gift from Drs. Eric Campeau & Paul Kaufman (Addgene plasmid # 17293; http://n2t.net/addgene:17293; RRID:Addgene_17293)(PMID: 19657394). psPAX2 and pMD2.G were gifts of the Duke Functional Genomics Facility. pcDNA3.1 myc-6xHis Sec24C was cloned as described (Cox et al., 2018). Myc-6xHis Sec24C was then subcloned into pENTR containing a multiple cloning site (gift from Dr. Vann Bennett) using Gibson Assembly (E5510, New England Biolabs). pENTR was linearized with NotI (R3189, New England Biolabs) and EcoRI (R3101, New England Biolabs). The following primers were designed using NEBuilder (http://nebuilder.neb.com/#!/):

F: 5’ – AAAGCAAGGCTCCGCGGCCGCCACCATGAAGCTTGCCACCATGGAA – 3’
R: 5’ – ATCCGCGGATCGTAGAATTCTTAGCTCAGTAGCTGCCG – 3’

Ser/Thr→Ala mutations were made in pENTR myc-6xHis Sec24C using methods previously described (Cox et al., 2018), but mutagenized DNA was transformed into One Shot Stbl3 Chemically Competent *E. coli* (C737303, ThermoFisher). Primers were designed using QuikChange Primer Design (http://www.genomics.agilent.com/primerDesignProgram.jsp). The following primers and their reverse complements were used:

S60A 5’ – CTCCCCAAGGTATGGCAAGAGCCCCACCT – 3’
S66A 5’ – AGGTGCCCCCGCGGAAGGTGGGG – 3’
S72A 5’ – GCCTGTGCTGTTGCGGCTGGAGGTGCC – 3’
S191A 5’ – CCTTGGGCCTGGGCAAGGGGAGGAGCCT – 3’
S773A 5’ – CTTTGGAGCTTTCTACATGGCCAACACGACAGATGTGGAG – 3’
T775A 5’ – CTTTCTACATGAGCAACGCGACAGATGTGGAGCTG – 3’
T776A 5’ – CTACATGAGCAACACGGCAGATGTGGAGCTGGC – 3’

Sec24D was subcloned from pEGFP Sec24D into pcDNA3.1 with an N-terminal myc-6xHis tag using Gibson Assembly (E5510, New England Biolabs). pcDNA3.1 was linearized with Not1 (R3189, New England Biolabs) and XbaI (R0145, New England Biolabs). Primers were designed using NEBuilder (http://nebuilder.neb.com/#!/). pEGFP-Sec24D was a gift from Dr. Henry Lester (Addgene plasmid # 32678; http://n2t.net/addgene:32678; RRID:Addgene_32678) (Richards et al., 2011). The following primers were used:

F: 5’ – ATCCAGCACAGTGGCGGCCGCAGTCAACAAGGTTACGTGG – 3’
R: 5’ – GGTTTAAACGGGCCCTCTAGATTAATTAAGCAGCTGACAGATC – 3’

Myc-6xHis Sec24D was then subcloned from pcDNA3.1 myc-6xHis Sec24D into pENTR containing a multiple cloning site (gift from Dr. Vann Bennett) using Gibson Assembly (E5510, New England Biolabs). pENTR was linearized with EcoRI (R3101, New England Biolabs) and AscI (R0558, New England Biolabs). Primers were designed using NEBuilder (http://nebuilder.neb.com/#!/). The following primers were used:

F: 5’ – TGCATCATCATCATCATCATGGAACAAAAACTCATCTCAGAAG – 3’
R: 5’ – AGAAAGCTGGGTCGGCTAATTAAGCAGCTGACAGATC – 3’

pENTR containing myc-6xHis Sec24C or Sec24D WT or mutants were used in a Gateway LR Clonase II Enzyme reaction (11791020, Invitrogen) according to manufacturer’s instructions with pLenti CMV/TO Puro DEST (670-1) (Addgene 1729).

pcDNA4-Sec31A-myc-6xHis and pcDNA4-eGFP-myc-6xHis were cloned as described previously (Cox et al., 2018). Sec31A Ser/Thr →Ala mutants were made in pcDNA4-Sec31A-myc-6xHis with primers designed at QuikChange Primer Design (http://www.genomics.agilent.com/primerDesignProgram.jsp). The following primers and their reverse complements were used:

S451A 5’ – CTTCAGCAGGCTGTGCAGGCACAAGGATTTATCAATTA – 3’
T658A 5’ – GCTTTAGCTGCAGTATTGGCTTATGCAAAGCCGGATG – 3’
S666A 5’ – ACTTATGCAAAGCCGGATGAATTTGCAGCCCTTTGTGA – 3’
T674A 5’ – AGCCCTTTGTGATCTTTTGGGAGCCAGGCTTGAAAv3’
S762A 5’ – GTTGGCAGCTCAGGGCGCTATTGCTGCAGCCTTG – 3’
T774A 5’ – GCTTTTCTTCCTGACAACGCCAACCAGCCAAATATCA – 3’
T1187A 5’ – AACTACTCAGAAGGATTGGCCATGCATACCCACATAG – 3’
T1190A 5’ – CTACTCAGAAGGATTGACCATGCATGCCCACATAGTTAG – 3’
S1202A 5’ – GCAACTTCAGTGAGACCGCTGCTTTCATGCCAGTT – 3’

Phusion polymerase (M0530, New England Biolabs) was utilized to perform mutagenesis reactions, after which samples were digested with DpnI, then transformed in One Shot TOP10 chemically competent *E. coli* (C404010, ThermoFisher) as described previously (Cox et al., 2018). Successful mutagenesis was confirmed by sequencing.

### Transfections

Cells were plated at ~20-50% confluence and transfected the following day using TransIT-293 transfection reagent (Mirus) and OPTI-MEM (1105802, ThermoFisher) as previously described (Cox et al., 2018).

### Immunoblotting (IB)

Samples were resolved on Tris-glycine SDS-PAGE gels and electroblotted onto PVDF (88518, ThermoFisher) or nitrocellulose membrane (162-115, BioRad) using standard methods. IBs were performed as previously described (Cox et al., 2018) and developed via enhanced chemiluminescence (ECL) (WesternBright ECL, Advansta) or infrared quantitative imaging using the Odyssey CLx (LI-COR) according to the manufacturer’s instructions. The following primary antibodies were used: rabbit anti-Sec23A (8162, Cell Signaling Technology; 1:2000), rabbit monoclonal anti-Sec24C (D9M4N) (14676S, Cell Signaling Technology; 1:2000), mouse monoclonal a-tubulin (T6074, Sigma-Aldrich; 1:100000), rabbit monoclonal GAPDH (14C10) (2188, Cell signaling Technology; 1:4000), mouse monoclonal anti-c-myc (9E10) (various vendors; 1:1000), mouse monoclonal anti-O-GlcNAc antibody 9D1 (various vendors; 1:1000), mouse monoclonal anti-O-GlcNAc 18B10.C7 (various vendors; 1:1000), mouse monoclonal anti-O-GlcNAc antibody 1F5 (various vendors; 1:1000), mouse monoclonal anti-O-GlcNAc antibody CTD110.6 (sc-59623, Santa Cruz Biotechnology; 1:1000), mouse monoclonal anti-O-GlcNAc antibody RL2 (SC-59624, Santa Cruz Biotech; 1:500), rabbit monoclonal anti-LC3A/B (D3U4C, Cell Signaling Technology; 1:1000), rabbit polyclonal anti-Sec31A (8506, Cell Signaling; 1:10000), rabbit monoclonal anti-Sec24D (D9M7L) (14687, Cell Signaling; 1:1000), mouse monoclonal anti-nucleoporin p62 (610498, BD Transduction Laboratories; 1:1000), and rabbit polyclonal anti-Sec13 (A303-980A-T, Bethyl Laboratories; 1:1000).

The following secondary antibodies were used: goat anti-mouse IgG (1030-05, horseradish peroxidase (HRP)-conjugated, SouthernBiotech; 1:10000), goat anti-rabbit IgG (4030-05, HRP-conjugated, SouthernBiotech; 1:10000), IRDye 800CW goat anti-mouse (925-32210, LI-COR Biosciences; 1:30000) and IRDye 800CW Goat anti-Rabbit IgG (H + L) (925-32211, LI-COR Biosciences; 1:30000).

### Immunoprecipitation (IP) and tandem purification

For myc IP, cells expressing myc-6xHis-tagged Sec24C, Sec24D or Sec31A were lysed in IP lysis buffer (1% Triton X-100, 150 mM NaCl, 1 mM EDTA, 20 mM Tris-HCl pH 7.4), with or without 0.1% SDS, supplemented with protease inhibitor cocktail (P8340, Sigma), 50 μM UDP (OGT inhibitor) and 5 μM PUGNAC (OGA inhibitor, 17151, Cayman Chemicals). Lysates were probe-sonicated, cleared by centrifugation, and quantified by BCA protein assay (23225, ThermoFisher) according to the manufacturer’s directions. IPs were performed on 2-10 mg total protein for IB or 50–100 mg total protein for MS analysis. Cleared lysates were adjusted to a final total protein concentration of ~2-5 mg/mL using IP lysis buffer supplemented with protease inhibitor, UDP and PUGNAc, and 1 μg (for myc-6xHis tagged Sec24) or 2ug (for myc-6xHis tagged Sec31) mouse monoclonal anti-c-myc antibody (9E10, various vendors) per 1 mg of total protein were added along with 30 μL washed protein A/G UltraLink Resin (53133, ThermoFisher) and rotated overnight at 4°C. Beads were washed three times with 1 mL of IP lysis buffer and then eluted. Elution was performed with IP lysis buffer plus 2X SDS-PAGE loading buffer (5X SDS-PAGE loading buffer: 250 mM Tris pH 6.8, 10% SDS, 30% glycerol, 5% β-mercaptoethanol, 0.02% bromophenol blue) and heating at 95 °C for 5 minutes. Eluents were then analyzed by SDS-PAGE and IB.

For tandem affinity purification, protein A/G beads were eluted twice in 500 μL using Ni-NTA wash buffer (8 M urea, 300 mM NaCl, 1% Triton X-100, 5 mM imidazole) with rotation. The two 500 μL elutions were pooled, and 30 μL washed 6xHisPur Ni-NTA resin (88223, ThermoFisher) was added to the eluate and the mixture was incubated at room temperature for 2 hours with end-over-end rotation. The Ni-NTA resin was washed three times with 1 mL of Ni-NTA wash buffer. The final elution from the Ni-NTA was performed using 8 M urea with 250 mM imidazole.

For endogenous Sec24C, Sec24D or Sec31A IP, cells were lysed in IP lysis buffer without SDS. The IP was then performed as above using 2-10 mg of protein and 1 μg of rabbit anti-Sec24C (A304-760A, Bethyl Laboratories) per 1 mg of protein or 0.9 μg of rabbit anti-Sec24D (14687, Cell Signaling Technology) or 1 μg of rabbit anti-Sec31A (Cell Signaling #8506) per 1 mg of protein.

### lonomycin treatment

HEK 293T cells stably expressing Sec31A-myc6xHIs were plated to be approximately 80% confluent the day of treatment. Ionomycin (I0634, Sigma-Aldrich) was added to the plate at a final concentration of 3 μM for 0-30 min. Cells were collected by scraping into cold phosphate-buffered saline (PBS) and centrifugation, lysed in IP lysis buffer with SDS, and analyzed by BCA assay, myc IP, SDS-PAGE and IB.

### OGT and OGA inhibition in live cells

OGT or OGA inhibition was achieved by incubating cells with fresh media containing 50 μM 5SG or 25-50 μM TG, respectively, for 24 hours at 37 °C in 5% CO_2_ prior to harvest, unless otherwise indicated.

### Rapamycin and tunicamycin treatment

For examination of Sec31 O-GlcNAcylation changes, HEK 293T cells were plated on 15 cm dishes and grown to ~80% confluency. Control samples were treated with DMSO, 50 μM TG, or 50 μM 5SG for 24 hours at 37 °C in 5% CO_2_. Experimental samples were treated in parallel with either 200 nM rapamycin for 16 hours or 2.5 μg/mL tunicamycin for 5 hours at 37 °C in 5% CO_2_. Cells were lysed, IP-ed with an anti-Sec31A antibody, and subjected to IBs. For Sec24C O-GlcNAcylation analysis, HEK 293T cells stably expressing myc-6xHis Sec24C were plated to be ~80% confluent the day of treatment. The next day, fresh media and 100 nM or 200 nM rapamycin or vehicle control were added to the plates. Cells were incubated at 37 °C in 5% CO_2_ for 16 hours. Cells were then lysed, a myc IP was performed, and samples were subjected to IBs.

### Generation of Sec24C^-/-^, Sec24D^-/-^ and Sec31A^-/-^ cells

CRISPR-Cas9 deletion of Sec24C, Sec24D and Sec31A from HEK 293T or HeLa cells was performed using transient expression of Cas9 and sgRNAs. Cells were transfected with pSpCas9(BB)-2A-GFP (PX458) + sgRNAs using methods previously described (Cox et al., 2018) and GFP positive cells were sorted into a 96-well plate, with each well containing a single cell, by the Duke Cancer Institute Flow Cytometry Core Facility using a BD Diva fluorescence-activated cell sorter. Surviving clones were expanded and Sec24C, Sec24D, or Sec31A deletion was verified by IB.

### Lentivirus generation

HEK 293T cells were plated to be 60-70% confluent in a 6 cm plate the following day. Cell culture medium was replaced prior to transfections. 150 μL pre-warmed OPTI-MEM was placed in 1.5 mL tubes with 6 μL TransIT-LT1 transfection reagent (MIR 2300, Mirus), vortexed briefly, and incubated for 15 minutes at room temperature. After incubation, 1 μg of pLenti CMV/TO Puro DEST (670-1) empty vector or containing myc-6xHis-tagged Sec24C or Sec24D WT or mutant, 900 ng of psPAX2, and 100 ng of pMD2.G were added to the 1.5 mL tubes, vortexed briefly and incubated for 15 minutes at room temperature. The transfection mixture was then added dropwise to the cells. Cells recovered at 37 °C and 5% CO_2_ for 12-18 hours. The media was then replaced with DMEM supplemented with 30% fetal bovine serum (FBS), 100 units/mL penicillin, and 100 μg/mL streptomycin, and cells were recovered at 37 °C and 5% CO_2_. 48 hours post-transfection, the medium containing virus was collected and stored at 4 °C. Fresh medium was added to the cells (DMEM supplemented with 30% FBS, 100 units/mL penicillin, and 100 μg/mL streptomycin) and the cells were recovered at 37 °C and 5% CO_2_. Virus was collected again 24 hours later. All virus was then centrifuged to pellet cellular debris, filtered through 45 μm low protein-binding filters and stored at −80 °C.

### Stable expression of myc-6xHis Sec24C wild type and unglycosylatable mutants

HEK 293T Sec24C^-/-^ or parental cells were plated in a 6-well plate to be ~50% confluent on the day of the infection. The medium was then changed and polybrene was added to a final concentration of 5 μg/mL. 1 mL of virus was added to each well. Plates were then centrifuged at 2000 rotations per minutes for 20 minutes. Cells were recovered at 37 °C and 5% CO_2_. 24-48 hours post-infection, cells were treated with puromycin (2 μg/mL for cell lines constructed in the parental HEK 293T background; 1 μg/mL for cell lines constructed in a single cell-derived clonal Sec24C’^7^’ background) until a mock-transfected plate no longer contained living cells.

### Stable expression of Sec31A-myc-6xHis wild type and unglycosylatable mutants

HEK 293T Sec31A^-/-^ cells were plated in 10-cm dishes, grown to 20% confluency and transfected with pcDNA4-eGFP-myc-6xHis, pcDNA4-Sec31A-myc-6xHis or pcDNA4-myc-6xHis expressing unglycosylatable Sec31A mutants. 36 hours post-transfection, 200 μg/mL Zeocin was added to the culture medium to select for successfully transfected cells. Cells were maintained in Zeocin-containing media until 100% of cells transfected with pcDNA4-eGFP-myc-6xHis were GFP-positive by fluorescence microscopy. Cells were grown and passaged one additional cycle, then subject to IB with Sec31A and myc antibodies to validate expression of a single myc-tagged Sec31A band at the expected molecular weight, with the eGFP-expressing line used as a negative control.

### Liquid chromatography–tandem MS (LC-MS/MS) analysis of Sec24C, Sec24D and Sec31A O-GlcNAcylation

Tandem-purified Sec24C, Sec24D or Sec31A was separated by SDS-PAGE and Coomassie stained. Stained bands of the correct molecular weight were subjected to standard in-gel trypsin digestion according to a Duke Proteomics and Metabolomics Shared Resource protocol (https://genome.duke.edu/sites/genome.duke.edu/files/In-gelDigestionProtocolrevised_0.pdf). Extracted peptides were lyophilized to dryness and resuspended in 12 μL of 0.2% formic acid/2% acetonitrile. Each sample was subjected to chromatographic separation on a Waters NanoAquity UPLC equipped with a 1.7 μm BEH130 C18 75 μm I.D. × 250 mm reversed-phase column. The mobile phase consisted of (A) 0.1% formic acid in water and (B) 0.1% formic acid in acetonitrile. Following a 4 μL injection, peptides were trapped for 3 minutes on a 5 μm Symmetry C18 180 μm I.D. × 20 mm column at 5 μL/min in 99.9% A. The analytical column was then switched in-line, and a linear elution gradient of 5% B to 40% B was performed over 60 minutes at 400 nL/min. The analytical column was connected to a fused silica PicoTip emitter (New Objective) with a 10 μm tip orifice and coupled to a Fusion Lumos Orbitrap mass spectrometer (Thermo Scientific) through an electrospray interface operating in data-dependent acquisition mode. The instrument was set to acquire a precursor MS scan from m/z 350 to 1800 every 3 seconds with a target AGC of 4e5. In data-dependent mode, MS/MS scans of the most abundant precursors were collected following higher-energy collisional dissociation (HCD) fragmentation at an HCD collision energy of 27%. Within the MS/MS spectra, if any diagnostic O-GlcNAc fragment ions (m/z 204.0867, 138.0545, or 366.1396) were observed, a second MS/MS spectrum of the precursor was acquired with electron transfer dissociation (ETD)/HCD fragmentation using charge-dependent ETD reaction times and either 30% or 15% supplemental collision energy for precursor charge states of at least +2. Target AGC settings were 3e5 and 250 ms at 30,000 resolution (at m/z 200). For all experiments, a 60-second dynamic exclusion was employed for previously fragmented precursor ions.

Raw LC-MS/MS data files were processed in Proteome Discoverer (Thermo Fisher Scientific) and then submitted to independent Mascot searches (Matrix Science) against a SwissProt database (human taxonomy) containing both forward and reverse entries of each protein (https://www.uniprot.org/proteomes/UP000005640) (20,322 forward entries). Search tolerances were 5 ppm for precursor ions and 0.02 Da for product ions using semi-trypsin specificity with up to two missed cleavages. Both *y/b*-type HCD and *c/z*-type ETD fragment ions were allowed for interpreting all spectra. Carbamidomethylation (+57.0214 Da on C) was set as a fixed modification, whereas oxidation (+15.9949 Da on M) and O-GlcNAc (+203.0794 Da on S/T) were considered dynamic mass modifications. All searched spectra were imported into Scaffold (v4.3, Proteome Software), and scoring thresholds were set to achieve a peptide false discovery rate of 1% using the PeptideProphet algorithm (http://peptideprophet.sourceforge.net/). When satisfactory ETD fragmentation was not obtained, HCD fragmentation was used to determine O-GlcNAc residue modification using the number of HexNAcs identified in combination with the number of serines and threonines in the peptide.

The mass spectrometry data for Sec24C, Sec24D and Sec31A O-GlcNAc site-mapping experiments have been deposited to the ProteomeXchange Consortium (Deutsch et al., 2017) via the MassIVE partner repository (https://massive.ucsd.edu/), with the dataset identifier MSV000086625. Reviewers may access complete raw data now via MassIVE using username “MSV000086625_reviewer” and password “a” (both without quotation marks). The data will be made public promptly upon publication.

## Supplemental Methods

### Quantitative RT-PCR (qRT-PCR)

Cells were plated in a 6-well plate to be 80% confluent the day of collection. Cells were lysed and RNA was extracted using the RNeasy Mini Kit (74104, Qiagen) according the manufacturer’s directions. The RTL buffer was supplemented with β-mercaptoethanol prior to lysis. RNA concentration was measured by Nanodrop 2000 Spectrophotometer (ND-2000, ThermoFisher Scientific) according to the manufacturer’s directions and RNA was stored at −80 °C. Reverse transcriptase reactions were performed using the High-Capacity cDNA Reverse Transcription Kit (4368814, ThermoFisher Scientific) according to the manufacturer’s directions. qRT-PCR reactions were performed on StepOnePlus Real-Time PCR System (Applied Biosystems) using Power SYBR™ Green PCR Master Mix (4367659, Applied Biosystems) and the following primers:

Sec24C F: 5’ – CAGACACAAGGGAAACAGAGA – 3’
Sec24C R: 5’ – GGGCAGGGATGTATGGAATAG – 3’
β-actin F: 5’ – CACTCTTCCAGCCTTCCTTC – 3’
β-actin R: 5’ – GTACAGGTCTTTGCGGATGT – 3’

Fold change in mRNA expression of Sec24C, compared to β-actin, was determined using the 2^-ΔΔCT^ method (Livak and Schmittgen, 2001).

### Ionomycin treatment

HEK 293T cells stably expressing myc-6xHis Sec24C were plated to be ~80% confluent the day of treatment. Ionomycin (I0634, Sigma Aldrich) was added to the plate at a final concentration of 1 μM (low treatment) or 3 μM (high treatment) for 1 hour. Cells were collected by scraping into PBS and centrifugation, lysed in IP lysis buffer without SDS, probe-sonicated, and analyzed by BCA assay, myc IP, SDS-PAGE and IB.

### Starvation-induced autophagy

HEK 293T or HeLa cells were maintained in Earle’s Balanced Salts Solution (EBSS) (E2888, Sigma Aldrich) for 1 to 5 hours or kept in DMEM as a negative control. Bafilomycin A (BAF) (B-1080, LC laboratories) was used at 100 nM for 1 hours. When EBSS and BAF were used together, cells were starved for 4 hours with EBSS and 100 nM BAF was added for one additional hour. Cells were collected by scraping into PBS and centrifugation, lysed in IP lysis buffer without SDS, and analyzed by BCA assay, Sec24C IP, SDS-PAGE and IB.

### Antibodies

Additional antibodies used were rabbit anti-TRAPα (gift of Dr. Christopher Nicchitta, Duke University, 1:8000), rabbit polyclonal anti-Sec24A (9678, Cell Signaling; 1:1000), and rabbit monoclonal Sec24B (D7D6S) (12042, Cell Signaling; 1:10000).

## Supplementary data

Figures S1–S3, Supplemental methods

## Funding

This work was supported by grant R01GM117473 from the National Institutes of Health to M.B.

## Acknowledgements

We thank members of the Boyce Lab for helpful discussion and advice, Dr. Erik Soderblom and the Duke Proteomics and Metabolomics Shared Resource for O-GlcNAc site-mapping analyses and Dr. John Lucocq for drawing our attention to early evidence linking COPII to O-GlcNAcylation.

## Conflict of interest statement

The authors declare that no conflicts of interest exist.

## Abbreviations

5SG: Ac45SGlcNAc
CID: collision-induced dissociation
COPII: coat protein II complex
CRISPR: clustered regularly interspaced short palindromic repeats
ER: endoplasmic reticulum
ETD: electron transfer dissociation
HBP: hexosamine biosynthetic pathway
IB: immunoblot
IP: immunoprecipitation
mTORC: mechanistic target of rapamycin complex
MS: mass spectrometry
O-GlcNAc: O-linked β-*N*-acetylglucosamine
OGA: O-GlcNAcase
OGT: O-GlcNAc transferase
PTMs: post-translational modifications
Rap: rapamycin
TG: Thiamet-G
UDP-GlcNAc: uridine diphosphate-GlcNAc

## Supplemental figure legends

**Figure S1.**
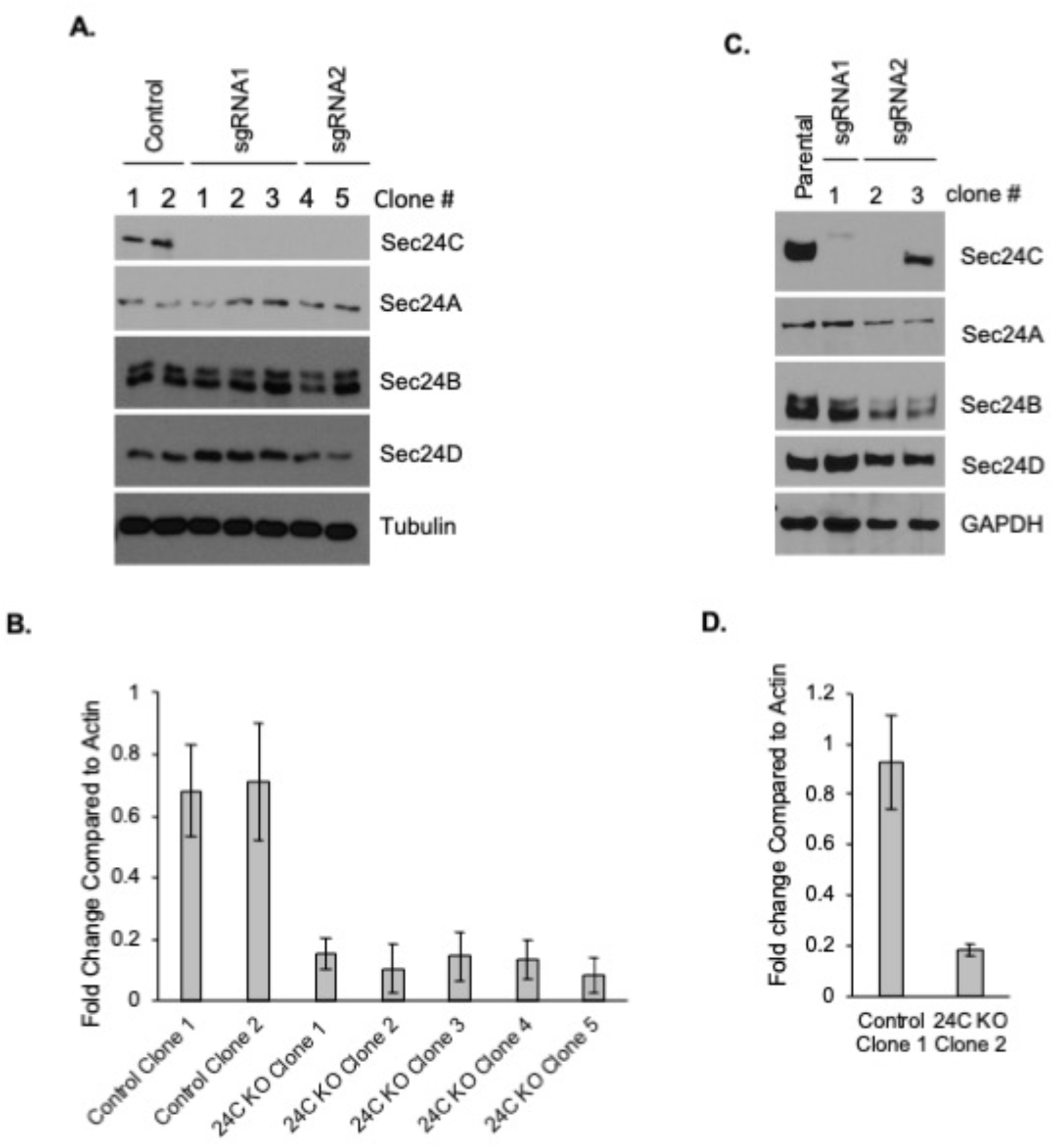
Validation of Sec24C^-/-^ HEK 293T and HeLa cells. **A.** HEK 293T cells were transfected with a construct expressing GFP, Cas9 and sgRNAs targeting Sec24C (two independent sgRNAs) or AAVS1 (control). GFP-positive cells were sorted by flow cytometry into a 96-well plate, such that each well contained a single cell. Successful deletion of Sec24C in single cell-derived clones was verified via IB. **B.** qRT-PCR verification of Sec24C^-/-^ clones (KO). Fold-change in Sec24C mRNA, compared to β-actin, was determined using the 2^-ΔΔCT^ method. **C.** Sec24C^-/-^ HeLa cells were derived as in **A.** and analyzed by IB. **D.** qRT-PCR verification of Sec24C^-/-^ HeLa cells as in **B.**

**Figure S2.**
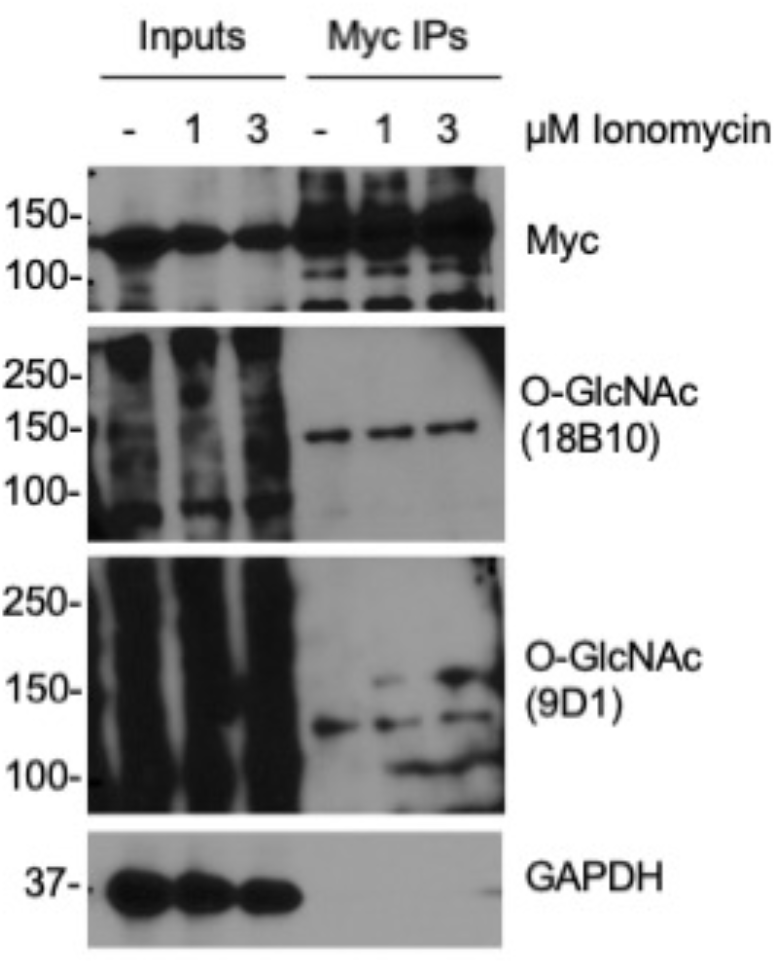
Ionomycin treatment has no discernible effect on Sec24C O-GlcNAcylation. HEK 293T cells stably expressing myc-6xHis-tagged Sec24C were treated with ionomycin as indicated for 1 hour. Sec24C was analyzed by myc IP and IB.

**Figure S3.**
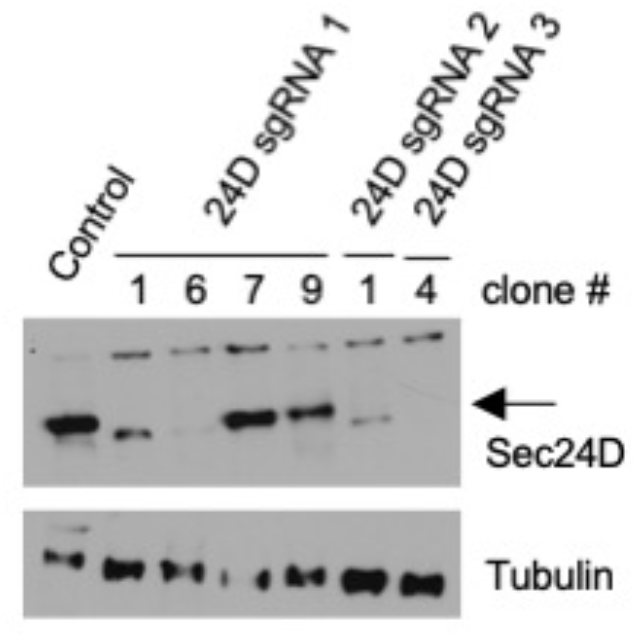
Validation of Sec24D^-/-^ cells. HEK 293T cells were transfected with a construct expressing GFP, Cas9 and sgRNAs against Sec24D (three independent sgRNAs) or AAVS1 (control). GFP-positive cells were sorted by flow cytometry into a 96-well plate, such that each well contained a single cell. Successful deletion of Sec24D in single cell-derived clones was verified via IB.

## Notes

https://massive.ucsd.edu/

### Competing Interest Statement

The authors have declared no competing interest.

## References

Adams, E.J., Chen, X.W., O’Shea, K.S., and Ginsburg, D. (2014). Mammalian COPII coat component SEC24C is required for embryonic development in mice. The Journal of Biological Chemistry 289, 20858.–20870.

Akan, I., Olivier-Van Stichelen, S., Bond, M.R., and Hanover, J.A. (2018). Nutrient-driven O-GlcNAc in proteostasis and neurodegeneration. Journal of neurochemistry 144, 7–34.

Al-Bari, M.A.A. (2020). A current view of molecular dissection in autophagy machinery. J Physiol Biochem.

Amodio, G., Renna, M., Paladino, S., Venturi, C., Tacchetti, C., Moltedo, O., Franceschelli, S., Mallardo, M., Bonatti, S., and Remondelli, P. (2009). Endoplasmic reticulum stress reduces the export from the ER and alters the architecture of post-ER compartments. The international journal of biochemistry & cell biology 41, 2511–2521.

Aridor, M. (2018). COPII gets in shape: Lessons derived from morphological aspects of early secretion. Traffic 19, 823–839.

Baines, A.C., Adams, E.J., Zhang, B., and Ginsburg, D. (2013). Disruption of the Sec24d gene results in early embryonic lethality in the mouse. PLoS ONE 8, e61114.

Barlowe, C. (2020). Twenty-five years after coat protein complex II. Molecular biology of the cell 31, 3–6.

Bethune, J., and Wieland, F.T. (2018). Assembly of COPI and COPII Vesicular Coat Proteins on Membranes. Annu Rev Biophys 47, 63–83.

Bhandari, D., Zhang, J., Menon, S., Lord, C., Chen, S., Helm, J.R., Thorsen, K., Corbett, K.D., Hay, J.C., and Ferro-Novick, S. (2013). Sit4p/PP6 regulates ER-to-Golgi traffic by controlling the dephosphorylation of COPII coat subunits. Molecular biology of the cell 24, 2727–2738.

Bhaskar, P.T., and Hay, N. (2007). The two TORCs and Akt. Developmental cell 12, 487–502.

Bi, X., Mancias, J.D., and Goldberg, J. (2007). Insights into COPII coat nucleation from the structure of Sec23.Sar1 complexed with the active fragment of Sec31. Dev Cell 13, 635–645.

Bond, M.R., and Hanover, J.A. (2013). O-GlcNAc cycling: a link between metabolism and chronic disease. Annual review of nutrition 33, 205–229.

Bond, M.R., and Hanover, J.A. (2015). A little sugar goes a long way: the cell biology of O-GlcNAc. The Journal of cell biology 208, 869–880.

Boyce, M., Carrico, I.S., Ganguli, A.S., Yu, S.-H., Hangauer, M.J., Hubbard, S.C., Kohler, J.J., and Bertozzi, C.R. (2011). Metabolic cross-talk allows labeling of O-linked beta-N-acetylglucosamine-modified proteins via the N-acetylgalactosamine salvage pathway. Proceedings of the National Academy of Sciences of the United States of America 108, 31413146.

Brandizzi, F. (2018). Transport from the endoplasmic reticulum to the Golgi in plants: Where are we now? Seminars in cell & developmental biology 80, 94–105.

Butkinaree, C., Park, K., and Hart, G.W. (2010). O-linked beta-N-acetylglucosamine (O-GlcNAc): Extensive crosstalk with phosphorylation to regulate signaling and transcription in response to nutrients and stress. Biochimica et biophysica acta 1800, 96–106.

Carlos Martín Zoppino, F., Damián Militello, R., Slavin, I., Álvarez, C., and Colombo, M.I. (2010). Autophagosome Formation Depends on the Small GTPase Rab1 and Functional ER Exit Sites. Traffic 11, 1246–1261.

Chen, X.W., Wang, H., Bajaj, K., Zhang, P., Meng, Z.X., Ma, D., Bai, Y., Liu, H.H., Adams, E., Baines, A., et al. (2013). SEC24A deficiency lowers plasma cholesterol through reduced PCSK9 secretion. Elife 2, e00444.

Cho, H.J., and Mook-Jung, I. (2018). O-GlcNAcylation regulates endoplasmic reticulum exit sites through Sec31A modification in conventional secretory pathway. FASEB J 32, 4641–4657.

Clark, P.M., Dweck, J.F., Mason, D.E., Hart, C.R., Buck, S.B., Peters, E.C., Agnew, B.J., and Hsieh-Wilson, L.C. (2008). Direct in-gel fluorescence detection and cellular imaging of O-GlcNAc-modified proteins. Journal of the American Chemical Society 130, 11576–11577.

Condon, K.J., and Sabatini, D.M. (2019). Nutrient regulation of mTORC1 at a glance. Journal of cell science 132.

Cox, N.J., Unlu, G., Bisnett, B.J., Meister, T.R., Condon, B.M., Luo, P.M., Smith, T.J., Hanna, M., Chhetri, A., Soderblom, E.J., et al. (2018). Dynamic Glycosylation Governs the Vertebrate COPII Protein Trafficking Pathway. Biochemistry 57, 91–107.

Cui, Y., Parashar, S., Zahoor, M., Needham, P.G., Mari, M., Zhu, M., Chen, S., Ho, H.C., Reggiori, F., Farhan, H., et al. (2019). A COPII subunit acts with an autophagy receptor to target endoplasmic reticulum for degradation. Science 365, 53–60.

Darley-Usmar, V.M., Ball, L.E., and Chatham, J.C. (2012). Protein O-linked beta-N-acetylglucosamine: a novel effector of cardiomyocyte metabolism and function. Journal of molecular and cellular cardiology 52, 538–549.

Dassanayaka, S., and Jones, S.P. (2014). O-GlcNAc and the cardiovascular system. Pharmacology & therapeutics 142, 62–71.

Davis, S., Wang, J., Zhu, M., Stahmer, K., Lakshminarayan, R., Ghassemian, M., Jiang, Y., Miller, E.A., and Ferro-Novick, S. (2016). Sec24 phosphorylation regulates autophagosome abundance during nutrient deprivation. eLife 5, e21167.

Dentin, R., Hedrick, S., Xie, J., Yates, J., 3rd, and Montminy, M. (2008). Hepatic glucose sensing via the CREB coactivator CRTC2. Science (New York, NY 319, 1402–1405.

Deutsch, E.W., Csordas, A., Sun, Z., Jarnuczak, A., Perez-Riverol, Y., Ternent, T., Campbell, D.S., Bernal-Llinares, M., Okuda, S., Kawano, S., et al. (2017). The ProteomeXchange consortium in 2017: supporting the cultural change in proteomics public data deposition. Nucleic Acids Res 45, D1100–D1106.

Dudognon, P., Maeder-Garavaglia, C., Carpentier, J.L., and Paccaud, J.P. (2004). Regulation of a COPII component by cytosolic O-glycosylation during mitosis. FEBS letters 561, 44–50.

Erickson, J.R. (2014). Mechanisms of CaMKII Activation in the Heart. Frontiers in pharmacology 5, 59.

Erickson, J.R., Pereira, L., Wang, L., Han, G., Ferguson, A., Dao, K., Copeland, R.J., Despa, F., Hart, G.W., Ripplinger, C.M., et al. (2013). Diabetic hyperglycaemia activates CaMKII and arrhythmias by O-linked glycosylation. Nature 502, 372–376.

Fahie, K., and Zachara, N.E. (2016). Molecular Functions of Glycoconjugates in Autophagy. Journal of molecular biology 428, 3305–3324.

Fath, S., Mancias, J.D., Bi, X.P., and Goldberg, J. (2007). Structure and organization of coat proteins in the COPII cage. Cell 129, 1325–1336.

Ge, L., Zhang, M., Kenny, S.J., Liu, D., Maeda, M., Saito, K., Mathur, A., Xu, K., and Schekman, R. (2017a). Remodeling of ER-exit sites initiates a membrane supply pathway for autophagosome biogenesis. EMBO Rep 18, 1586–1603.

Ge, L., Zhang, M., Kenny, S.J., Liu, D., Maeda, M., Saito, K., Mathur, A., Xu, K., and Schekman, R. (2017b). Remodeling of ER-exit sites initiates a membrane supply pathway for autophagosome biogenesis. EMBO reports 18, 1586–1603.

Ge, L., Zhang, M., and Schekman, R. (2014). Phosphatidylinositol 3-kinase and COPII generate LC3 lipidation vesicles from the ER-Golgi intermediate compartment. eLife 3, e04135–e04135.

Gloster, T.M., Zandberg, W.F., Heinonen, J.E., Shen, D.L., Deng, L., and Vocadlo, D.J. (2011). Hijacking a biosynthetic pathway yields a glycosyltransferase inhibitor within cells. Nature chemical biology 7, 174–181.

Gomez-Navarro, N., and Miller, E.A. (2016). COP-coated vesicles. Curr Biol 26, R54–R57.

Graef, M., Friedman, J.R., Graham, C., Babu, M., and Nunnari, J. (2013). ER exit sites are physical and functional core autophagosome biogenesis components. Molecular Biology of the Cell 24, 2918–2931.

Han, I., and Kudlow, J.E. (1997). Reduced O glycosylation of Sp1 is associated with increased proteasome susceptibility. Molecular and cellular biology 17, 2550–2558.

Hanover, J.A., Krause, M.W., and Love, D.C. (2010). The hexosamine signaling pathway: O-GlcNAc cycling in feast or famine. Biochimica et biophysica acta 1800, 80–95.

Hanover, J.A., and Wang, P. (2013). O-GlcNAc cycling shows neuroprotective potential in C. elegans models of neurodegenerative disease. Worm 2, e27043.

Hardiville, S., and Hart, G.W. (2014). Nutrient regulation of signaling, transcription, and cell physiology by O-GlcNAcylation. Cell metabolism 20, 208–213.

Hart, G.W. (2014). Three Decades of Research on O-GlcNAcylation - A Major Nutrient Sensor That Regulates Signaling, Transcription and Cellular Metabolism. Frontiers in endocrinology 5, 183.

Hart, G.W., Slawson, C., Ramirez-Correa, G., and Lagerlof, O. (2011). Cross talk between O-GlcNAcylation and phosphorylation: roles in signaling, transcription, and chronic disease. Annual review of biochemistry 80, 825–858.

Helm, J.R., Bentley, M., Thorsen, K.D., Wang, T., Foltz, L., Oorschot, V., Klumperman, J., and Hay, J.C. (2014). Apoptosis-linked gene-2 (ALG-2)/Sec31 interactions regulate endoplasmic reticulum (ER)-to-Golgi transport: a potential effector pathway for luminal calcium. The Journal of Biological Chemistry 289, 23609–23628.

Hu, H., Gourguechon, S., Wang, C.C., and Li, Z. (2016). The G1 Cyclin-dependent Kinase CRK1 in Trypanosoma brucei Regulates Anterograde Protein Transport by Phosphorylating the COPII Subunit Sec31. The Journal of Biological Chemistry 291, 15527–15539.

Hutchings, J., Stancheva, V., Miller, E.A., and Zanetti, G. (2018). Subtomogram averaging of COPII assemblies reveals how coat organization dictates membrane shape. Nat Commun 9, 4154.

Hutchings, J., and Zanetti, G. (2019). Coat flexibility in the secretory pathway: a role in transport of bulky cargoes. Current opinion in cell biology 59, 104–111.

Ishihara, N., Hamasaki, M., Yokota, S., Suzuki, K., Kamada, Y., Kihara, A., Yoshimori, T., Noda, T., and Ohsumi, Y. (2001). Autophagosome requires specific early Sec proteins for its formation and NSF/SNARE for vacuolar fusion. Molecular biology of the cell 12, 3690–3702.

Jeong, Y.T., Simoneschi, D., Keegan, S., Melville, D., Adler, N.S., Saraf, A., Florens, L., Washburn, M.P., Cavasotto, C.N., Fenyo, D., et al. (2018). The ULK1-FBXW5-SEC23B nexus controls autophagy. Elife 7.

Jin, L., Pahuja, K.B., Wickliffe, K.E., Gorur, A., Baumgartel, C., Schekman, R., and Rape, M. (2012). Ubiquitin-dependent regulation of COPII coat size and function. Nature 482, 495–500.

Karanasios, E., Walker, S.A., Okkenhaug, H., Manifava, M., Hummel, E., Zimmermann, H., Ahmed, Q., Domart, M.-C., Collinson, L., and Ktistakis, N.T. (2016). Autophagy initiation by ULK complex assembly on ER tubulovesicular regions marked by ATG9 vesicles. Nature communications 7, 12420–12420.

Keembiyehetty, C., Love, D.C., Harwood, K.R., Gavrilova, O., Comly, M.E., and Hanover, J.A. (2015). Conditional knockout reveals a requirement for O-GlcNAcase in metabolic homeostasis. The Journal of Biological Chemistry.

Kettenbach, A.N., Schweppe, D.K., Faherty, B.K., Pechenick, D., Pletnev, A.A., and Gerber, S.A. (2011). Quantitative Phosphoproteomics Identifies Substrates and Functional Modules of Aurora and Polo-Like Kinase Activities in Mitotic Cells. Science signaling 4, 10.1126/scisignal.2001497.

Kim, J., and Guan, K.L. (2019). mTOR as a central hub of nutrient signalling and cell growth. Nature cell biology 21, 63–71.

King, D.T., Males, A., Davies, G.J., and Vocadlo, D.J. (2019). Molecular mechanisms regulating O-linked N-acetylglucosamine (O-GlcNAc)-processing enzymes. Current opinion in chemical biology 53, 131–144.

Koreishi, M., Yu, S., Oda, M., Honjo, Y., and Satoh, A. (2013). CK2 phosphorylates Sec31 and regulates ER-To-Golgi trafficking. PLoS ONE 8, e54382.

la Cour, J.M., Schindler, A.J., Berchtold, M.W., and Schekman, R. (2013). ALG-2 attenuates COPII budding in vitro and stabilizes the Sec23/Sec31A complex. PLoS ONE 8, e75309.

Lazarus, B.D., Love, D.C., and Hanover, J.A. (2009). O-GlcNAc cycling: implications for neurodegenerative disorders. The international journal of biochemistry & cell biology 41, 2134–2146.

Lederkremer, G.Z., Cheng, Y., Petre, B.M., Vogan, E., Springer, S., Schekman, R., Walz, T., and Kirchhausen, T. (2001). Structure of the Sec23p/24p and Sec13p/31p complexes of COPII. Proc Natl Acad Sci U S A 98, 10704–10709.

Lee, A., Miller, D., Henry, R., Paruchuri, V.D., O’Meally, R.N., Boronina, T., Cole, R.N., and Zachara, N.E. (2016). Combined Antibody/Lectin Enrichment Identifies Extensive Changes in the O-GlcNAc Sub-proteome upon Oxidative Stress. Journal of proteome research.

Leney, A.C., El Atmioui, D., Wu, W., Ovaa, H., and Heck, A.J.R. (2017). Elucidating crosstalk mechanisms between phosphorylation and O-GlcNAcylation. Proceedings of the National Academy of Sciences of the United States of America.

Levine, Z.G., and Walker, S. (2016). The Biochemistry of O-GlcNAc Transferase: Which Functions Make It Essential in Mammalian Cells? Annual review of biochemistry 85, 631–657.

Liu, L., Cai, J., Wang, H., Liang, X., Zhou, Q., Ding, C., Zhu, Y., Fu, T., Guo, Q., Xu, Z., et al. (2019). Coupling of COPII vesicle trafficking to nutrient availability by the IRE1alpha-XBP1s axis. Proceedings of the National Academy of Sciences of the United States of America 116, 11776–11785.

Livak, K.J., and Schmittgen, T.D. (2001). Analysis of relative gene expression data using realtime quantitative PCR and the 2(-Delta Delta C(T)) Method. Methods 25, 402–408.

Lord, C., Bhandari, D., Menon, S., Ghassemian, M., Nycz, D., Hay, J., Ghosh, P., and Ferro-Novick, S. (2011). Sequential interactions with Sec23 control the direction of vesicle traffic. Nature 473, 181–186.

Lucocq, J.M., Berger, E.G., and Roth, J. (1987). Detection of terminal N-linked N-acetylglucosamine residues in the Golgi apparatus using galactosyltransferase and endoglucosaminidase F/peptide N-glycosidase F: adaptation of a biochemical approach to electron microscopy. J Histochem Cytochem 35, 67–74.

Ma, J., and Hart, G.W. (2013). Protein O-GlcNAcylation in diabetes and diabetic complications. Expert review of proteomics 10, 365–380.

Ma, Z., and Vosseller, K. (2013). O-GlcNAc in cancer biology. Amino acids 45, 719–733.

Markova, E.A., and Zanetti, G. (2019). Visualizing membrane trafficking through the electron microscope: cryo-tomography of coat complexes. Acta Crystallogr D Struct Biol 75, 467–474.

Matsuoka, K., Orci, L., Amherdt, M., Bednarek, S.Y., Hamamoto, S., Schekman, R., and Yeung, T. (1998). COPII-coated vesicle formation reconstituted with purified coat proteins and chemically defined liposomes. Cell 93, 263–275.

Matsuoka, K., Schekman, R., Orci, L., and Heuser, J.E. (2001). Surface structure of the COPII-coated vesicle. Proc Natl Acad Sci U S A 98, 13705–13709.

McGourty, C.A., Akopian, D., Walsh, C., Gorur, A., Werner, A., Schekman, R., Bautista, D., and Rape, M. (2016). Regulation of the CUL3 Ubiquitin Ligase by a Calcium-Dependent Co-adaptor. Cell 167, 525–538 e514.

Merte, J., Jensen, D., Wright, K., Sarsfield, S., Wang, Y., Schekman, R., and Ginty, D.D. (2010). Sec24b selectively sorts Vangl2 to regulate planar cell polarity during neural tube closure. Nature cell biology 12, 41–46; sup pp 41-48.

Myers, S.A., Daou, S., Affar el, B., and Burlingame, A. (2013). Electron transfer dissociation (ETD): the mass spectrometric breakthrough essential for O-GlcNAc protein site assignments-a study of the O-GlcNAcylated protein host cell factor C1. Proteomics 13, 982–991.

Ong, Y.S., Tang, B.L., Loo, L.S., and Hong, W. (2010). p125A exists as part of the mammalian Sec13/Sec31 COPII subcomplex to facilitate ER-Golgi transport. The Journal of cell biology 190, 331–345.

Peotter, J., Kasberg, W., Pustova, I., and Audhya, A. (2019). COPII-mediated trafficking at the ER/ERGIC interface. Traffic 20, 491–503.

Pravata, V.M., Gundogdu, M., Bartual, S.G., Ferenbach, A.T., Stavridis, M., Ounap, K., Pajusalu, S., Zordania, R., Wojcik, M.H., and van Aalten, D.M.F. (2020a). A missense mutation in the catalytic domain of O-GlcNAc transferase links perturbations in protein O-GlcNAcylation to X-linked intellectual disability. FEBS letters 594, 717–727.

Pravata, V.M., Muha, V., Gundogdu, M., Ferenbach, A.T., Kakade, P.S., Vandadi, V., Wilmes, A.C., Borodkin, V.S., Joss, S., Stavridis, M.P., et al. (2019). Catalytic deficiency of O-GlcNAc transferase leads to X-linked intellectual disability. Proceedings of the National Academy of Sciences of the United States of America 116, 14961–14970.

Pravata, V.M., Omelkova, M., Stavridis, M.P., Desbiens, C.M., Stephen, H.M., Lefeber, D.J., Gecz, J., Gundogdu, M., Ounap, K., Joss, S., et al. (2020b). An intellectual disability syndrome with single-nucleotide variants in O-GlcNAc transferase. Eur J Hum Genet 28, 706–714.

Ran, F.A., Hsu, P.D., Wright, J., Agarwala, V., Scott, D.A., and Zhang, F. (2013). Genome engineering using the CRISPR-Cas9 system. Nature protocols 8, 2281–2308.

Richards, C.I., Srinivasan, R., Xiao, C., Mackey, E.D., Miwa, J.M., and Lester, H.A. (2011). Trafficking of alpha4* nicotinic receptors revealed by superecliptic phluorin: effects of a beta4 amyotrophic lateral sclerosis-associated mutation and chronic exposure to nicotine. The Journal of Biological Chemistry 286, 31241–31249.

Ruan, H.B., Ma, Y., Torres, S., Zhang, B., Feriod, C., Heck, R.M., Qian, K., Fu, M., Li, X., Nathanson, M.H., et al. (2017). Calcium-dependent O-GlcNAc signaling drives liver autophagy in adaptation to starvation. Genes & development 31, 1655–1665.

Sadelain, M., Papapetrou, E.P., and Bushman, F.D. (2011). Safe harbours for the integration of new DNA in the human genome. Nature reviews Cancer 12, 51–58.

Salama, N.R., Chuang, J.S., and Schekman, R.W. (1997). Sec31 encodes an essential component of the COPII coat required for transport vesicle budding from the endoplasmic reticulum. Molecular biology of the cell 8, 205–217.

Sarmah, S., Barrallo-Gimeno, A., Melville, D.B., Topczewski, J., Solnica-Krezel, L., and Knapik, E.W. (2010). Sec24D-dependent transport of extracellular matrix proteins is required for zebrafish skeletal morphogenesis. PLoS ONE 5, e10367.

Saxton, R.A., and Sabatini, D.M. (2017). mTOR Signaling in Growth, Metabolism, and Disease. Cell 168, 960–976.

Selvan, N., George, S., Serajee, F.J., Shaw, M., Hobson, L., Kalscheuer, V., Prasad, N., Levy, S.E., Taylor, J., Aftimos, S., et al. (2018). O-GlcNAc transferase missense mutations linked to X-linked intellectual disability deregulate genes involved in cell fate determination and signaling. The Journal of Biological Chemistry 293, 10810–10824.

Shafi, R., Iyer, S.P., Ellies, L.G., O’Donnell, N., Marek, K.W., Chui, D., Hart, G.W., and Marth, J.D. (2000). The O-GlcNAc transferase gene resides on the X chromosome and is essential for embryonic stem cell viability and mouse ontogeny. Proceedings of the National Academy of Sciences of the United States of America 97, 5735–5739.

Sharpe, L.J., Luu, W., and Brown, A.J. (2011). Akt phosphorylates Sec24: new clues into the regulation of ER-to-Golgi trafficking. Traffic 12, 19–27.

Shin, S.H., Love, D.C., and Hanover, J.A. (2011). Elevated O-GlcNAc-dependent signaling through inducible mOGT expression selectively triggers apoptosis. Amino acids 40, 885–893.

Singh, J.P., Zhang, K., Wu, J., and Yang, X. (2015). O-GlcNAc signaling in cancer metabolism and epigenetics. Cancer letters 356, 244–250.

Stagg, S.M., Gurkan, C., Fowler, D.M., LaPointe, P., Foss, T.R., Potter, C.S., Carragher, B., and Balch, W.E. (2006). Structure of the Sec13/31 COPII coat cage. Nature 439, 234–238.

Stagg, S.M., LaPointe, P., Razvi, A., Gurkan, C., Potter, C.S., Carragher, B., and Balch, W.E. (2008). Structural basis for cargo regulation of COPII coat assembly. Cell 134, 474–484.

Stancheva, V.G., Hutchings, J., Li, X.-H., Santhanam, B., Babu, M.M., Zanetti, G., and Miller, E.A. (2020). A multivalent fuzzy interface drives reversible COPII coat assembly. bioRxiv, 2020.2004.2015.043356.

Tarbet, H.J., Dolat, L., Smith, T.J., Condon, B.M., O’Brien, E.T., 3rd, Valdivia, R.H., and Boyce, M. (2018a). Site-specific glycosylation regulates the form and function of the intermediate filament cytoskeleton. Elife 7.

Tarbet, H.J., Toleman, C.A., and Boyce, M. (2018b). A Sweet Embrace: Control of ProteinProtein Interactions by O-Linked beta-N-Acetylglucosamine. Biochemistry 57, 13–21.

Tarrant, M.K., Rho, H.S., Xie, Z., Jiang, Y.L., Gross, C., Culhane, J.C., Yan, G., Qian, J., Ichikawa, Y., Matsuoka, T., et al. (2012). Regulation of CK2 by phosphorylation and O-GlcNAcylation revealed by semisynthesis. Nature chemical biology 8, 262–269.

Taylor, R.P., Geisler, T.S., Chambers, J.H., and McClain, D.A. (2009). Up-regulation of O-GlcNAc transferase with glucose deprivation in HepG2 cells is mediated by decreased hexosamine pathway flux. The Journal of Biological Chemistry 284, 3425–3432.

Teo, C.F., Ingale, S., Wolfert, M.A., Elsayed, G.A., Not, L.G., Chatham, J.C., Wells, L., and Boons, G.J. (2010). Glycopeptide-specific monoclonal antibodies suggest new roles for O-GlcNAc. Nature chemical biology 6, 338–343.

Toleman, C.A., Schumacher, M.A., Yu, S.H., Zeng, W., Cox, N.J., Smith, T.J., Soderblom, E.J., Wands, A.M., Kohler, J.J., and Boyce, M. (2018). Structural basis of O-GlcNAc recognition by mammalian 14-3-3 proteins. Proceedings of the National Academy of Sciences of the United States of America 115, 5956–5961.

Vaidyanathan, K., Durning, S., and Wells, L. (2014). Functional O-GlcNAc modifications: implications in molecular regulation and pathophysiology. Critical reviews in biochemistry and molecular biology 49, 140–163.

Vaidyanathan, K., Niranjan, T., Selvan, N., Teo, C.F., May, M., Patel, S., Weatherly, B., Skinner, C., Opitz, J., Carey, J., et al. (2017). Identification and characterization of a missense mutation in the O-linked beta-N-acetylglucosamine (O-GlcNAc) transferase gene that segregates with X-linked intellectual disability. The Journal of Biological Chemistry 292, 8948–8963.

Vaidyanathan, K., and Wells, L. (2014). Multiple tissue-specific roles for the O-GlcNAc post-translational modification in the induction of and complications arising from type II diabetes. The Journal of Biological Chemistry 289, 34466–34471.

Vercoutter-Edouart, A.S., Yazidi-Belkoura, I.E., Guinez, C., Baldini, S., Leturcq, M., Mortuaire, M., Mir, A.M., Steenackers, A., Dehennaut, V., Pierce, A., et al. (2015). Detection and identification of O-GlcNAcylated proteins by proteomic approaches. Proteomics 15, 1039–1050.

Wang, B., Joo, J.H., Mount, R., Teubner, B.J.W., Krenzer, A., Ward, A.L., Ichhaporia, V.P., Adams, E.J., Khoriaty, R., Peters, S.T., et al. (2018). The COPII cargo adapter SEC24C is essential for neuronal homeostasis. The Journal of clinical investigation 128, 3319–3332.

Wang, P., and Hanover, J.A. (2013). Nutrient-driven O-GlcNAc cycling influences autophagic flux and neurodegenerative proteotoxicity. Autophagy 9, 604–606.

Wang, S., Huang, X., Sun, D., Xin, X., Pan, Q., Peng, S., Liang, Z., Luo, C., Yang, Y., Jiang, H., et al. (2012). Extensive crosstalk between O-GlcNAcylation and phosphorylation regulates Akt signaling. PLoS ONE 7, e37427.

Wansleeben, C., Feitsma, H., Montcouquiol, M., Kroon, C., Cuppen, E., and Meijlink, F. (2010). Planar cell polarity defects and defective Vangl2 trafficking in mutants for the COPII gene Sec24b. Development 137, 1067–1073.

Wells, L., Vosseller, K., Cole, R.N., Cronshaw, J.M., Matunis, M.J., and Hart, G.W. (2002). Mapping sites of O-GlcNAc modification using affinity tags for serine and threonine post-translational modifications. Mol Cell Proteomics 1, 791–804.

Willems, A.P., Gundogdu, M., Kempers, M.J.E., Giltay, J.C., Pfundt, R., Elferink, M., Loza, B.F., Fuijkschot, J., Ferenbach, A.T., van Gassen, K.L.I., et al. (2017). Mutations in N-acetylglucosamine (O-GlcNAc) transferase in patients with X-linked intellectual disability. The Journal of Biological Chemistry 292, 12621–12631.

Woo, C.M., Iavarone, A.T., Spiciarich, D.R., Palaniappan, K.K., and Bertozzi, C.R. (2015). Isotope-targeted glycoproteomics (IsoTaG): a mass-independent platform for intact N-and O-glycopeptide discovery and analysis. Nature methods 12, 561–567.

Woo, C.M., Lund, P.J., Huang, A.C., Davis, M.M., Bertozzi, C.R., and Pitteri, S.J. (2018). Mapping and Quantification of Over 2000 O-linked Glycopeptides in Activated Human T Cells with Isotope-Targeted Glycoproteomics (Isotag). Mol Cell Proteomics 17, 764–775.

Yamasaki, A., Tani, K., Yamamoto, A., Kitamura, N., and Komada, M. (2006). The Ca2+-binding protein ALG-2 is recruited to endoplasmic reticulum exit sites by Sec31A and stabilizes the localization of Sec31A. Mol Biol Cell 17, 4876–4887.

Yang, X., and Qian, K. (2017). Protein O-GlcNAcylation: emerging mechanisms and functions. Nature reviews Molecular cell biology.

Yang, Y.R., Song, M., Lee, H., Jeon, Y., Choi, E.J., Jang, H.J., Moon, H.Y., Byun, H.Y., Kim, E.K., Kim, D.H., et al. (2012). O-GlcNAcase is essential for embryonic development and maintenance of genomic stability. Aging cell 11, 439–448.

Yi, W., Clark, P.M., Mason, D.E., Keenan, M.C., Hill, C., Goddard, W.A., 3rd, Peters, E.C., Driggers, E.M., and Hsieh-Wilson, L.C. (2012). Phosphofructokinase 1 glycosylation regulates cell growth and metabolism. Science (New York, NY 337, 975–980.

Yuzwa, S.A., Macauley, M.S., Heinonen, J.E., Shan, X., Dennis, R.J., He, Y., Whitworth, G.E., Stubbs, K.A., McEachern, E.J., Davies, G.J., et al. (2008). A potent mechanism-inspired O-GlcNAcase inhibitor that blocks phosphorylation of tau in vivo. Nature chemical biology 4, 483490.

Yuzwa, S.A., Shan, X., Macauley, M.S., Clark, T., Skorobogatko, Y., Vosseller, K., and Vocadlo, D.J. (2012). Increasing O-GlcNAc slows neurodegeneration and stabilizes tau against aggregation. Nature chemical biology 8, 393–399.

Yuzwa, S.A., and Vocadlo, D.J. (2014). O-GlcNAc and neurodegeneration: biochemical mechanisms and potential roles in Alzheimer’s disease and beyond. Chemical Society reviews 43, 6839–6858.

Zachara, N.E., Molina, H., Wong, K.Y., Pandey, A., and Hart, G.W. (2011). The dynamic stress-induced “O-GlcNAc-ome” highlights functions for O-GlcNAc in regulating DNA damage/repair and other cellular pathways. Amino acids 40, 793–808.

Zachara, N.E., O’Donnell, N., Cheung, W.D., Mercer, J.J., Marth, J.D., and Hart, G.W. (2004). Dynamic O-GlcNAc modification of nucleocytoplasmic proteins in response to stress. A survival response of mammalian cells. The Journal of Biological Chemistry 279, 30133–30142.

Zacharogianni, M., Aguilera-Gomez, A., Veenendaal, T., Smout, J., and Rabouille, C. (2014). A stress assembly that confers cell viability by preserving ERES components during amino-acid starvation. Elife 3.

Zhang, F., Su, K., Yang, X., Bowe, D.B., Paterson, A.J., and Kudlow, J.E. (2003). O-GlcNAc modification is an endogenous inhibitor of the proteasome. Cell 115, 715–725.

Zhong, J., Martinez, M., Sengupta, S., Lee, A., Wu, X., Chaerkady, R., Chatterjee, A., O’Meally, R.N., Cole, R.N., Pandey, A., et al. (2015). Quantitative phosphoproteomics reveals crosstalk between phosphorylation and O-GlcNAc in the DNA damage response pathway. Proteomics 15, 591–607.

Zhu, Y., Shan, X., Yuzwa, S.A., and Vocadlo, D.J. (2014). The emerging link between O-GlcNAc and Alzheimer disease. The Journal of Biological Chemistry 289, 34472–34481.

Zou, L., Zhu-Mauldin, X., Marchase, R.B., Paterson, A.J., Liu, J., Yang, Q., and Chatham, J.C. (2012). Glucose deprivation-induced increase in protein O-GlcNAcylation in cardiomyocytes is calcium-dependent. The Journal of Biological Chemistry 287, 34419–34431.

